# Diet-induced phospholipid remodeling dictates ferroptosis sensitivity and tumorigenesis in the pancreas

**DOI:** 10.1101/2025.04.04.645324

**Authors:** Christian Felipe Ruiz, Xiangyu Ge, Rylee McDonnell, Sherry S. Agabiti, Daniel C. McQuaid, Andy Tang, Meera Kharwa, Jennifer Goodell, Rocío del M. Saavedra-Peña, Allison Wing, Guangtao Li, Natasha Pinto Medici, Marie E. Robert, Rohan R. Varshney, Michael C. Rudolph, Fred S. Gorelick, John Wysolmerski, Daniel Canals, John D. Haley, Matthew S. Rodeheffer, Mandar Deepak Muzumdar

**Affiliations:** Department of Genetics, Yale University School of Medicine; New Haven, CT 06520, USA; Yale Cancer Biology Institute, Yale University; West Haven, CT 06516, USA; Translational Molecular Medicine, Pharmacology, and Physiology Program, Yale University; New Haven, CT 06510, USA; M.D.-Ph.D. Program, Yale University; New Haven, CT 06510, USA; Department of Comparative Medicine, Yale University School of Medicine; New Haven, CT 06520, USA; Department of Molecular, Cellular, and Developmental Biology, Yale University; New Haven, CT 06520, USA; Department of Biochemistry and Cell Biology, Stony Brook University; Stony Brook, NY 11794, USA; Department of Therapeutic Radiology, Yale University School of Medicine; New Haven, CT 06510, USA; Department of Pathology, Yale University School of Medicine; New Haven, CT 06510, USA; Yale Cancer Center and Smilow Cancer Hospital, Yale University, New Haven, CT; Department of Biochemistry and Physiology and Harold Hamm Diabetes Center, University of Oklahoma University Health Sciences; Oklahoma City, OK 73104, USA; Department of Internal Medicine, Section of Digestive Diseases, Yale University School of Medicine; New Haven, CT 06510, USA; Department of Cell Biology, Yale University School of Medicine; New Haven, CT 06510, USA; Veterans Affairs Health Care System, Yale University; New Haven, CT 06510, USA; Department of Internal Medicine, Section of Endocrinology, Yale University School of Medicine; New Haven, CT 06510, USA; Department of Medicine, Renaissance School of Medicine, Stony Brook University; Stony Brook, NY 11794, USA; Department of Pathology, Renaissance School of Medicine, Stony Brook University; Stony Brook, NY 11794, USA; Department of Cellular and Molecular Physiology, Yale University School of Medicine, New Haven, CT 06520, USA; Department of Internal Medicine, Section of Medical Oncology, Yale University School of Medicine; New Haven, CT 06510, USA; Molecular Cell Biology, Genetics, and Development Program, Yale University; New Haven, CT 06510, USA

**Keywords:** pancreatic cancer, fatty acid metabolism, diet, obesity, ferroptosis

## Abstract

High-fat diet (HFD) intake has been linked to an increased risk of pancreatic ductal adenocarcinoma (PDAC), a lethal and therapy-resistant cancer. However, whether and how specific dietary fats drive cancer development remains unresolved. Leveraging an oncogenic *Kras*-driven mouse model that closely mimics human PDAC progression, we screened a dozen isocaloric HFDs differing solely in fat source and representing the diversity of human fat consumption. Unexpectedly, diets rich in oleic acid – a monounsaturated fatty acid (MUFA) typically associated with good health – markedly enhanced tumorigenesis. Conversely, diets high in polyunsaturated fatty acids (PUFAs) suppressed tumor progression. Relative dietary fatty acid saturation levels (PUFA/MUFA) governed pancreatic membrane phospholipid composition, lipid peroxidation, and ferroptosis sensitivity in mice, concordant with circulating PUFA/MUFA levels being linked to altered PDAC risk in humans. These findings directly implicate dietary unsaturated fatty acids in controlling ferroptosis susceptibility and tumorigenesis, supporting potential “precision nutrition” strategies for PDAC prevention.

## INTRODUCTION

Pancreatic ductal adenocarcinoma (PDAC) remains one of the most deadly cancers, with a 5-year survival rate of only ∼13%^1^. Obesity and diet have emerged as key modifiable risk factors, with epidemiological studies demonstrating a positive correlation between body-mass index (BMI) or fat consumption and PDAC risk^2–5^. Per-capita fat intake has risen rapidly worldwide, and this increase is thought to be a major contributor to growing PDAC incidence^6^. Consistent with these observations, preclinical studies have shown that high-fat diet (HFD) feeding promotes tumor progression in transplant and autochthonous PDAC mouse models^7–12^. However, research linking HFD to cancer has been limited by numerous confounding variables, including: (1) variability in the percentage of diet derived from fat (30-60% of kilocalories (kcal)); (2) treatment duration (weeks to months); specific models used (transplant vs. autochthonous); (4) mouse strain; (5) sex; and (6) fat source^12–14^. Most cancer studies to date have used a 60% kcal lard HFD, which surpasses even the highest quartile of human fat intake^15^. Furthermore, this diet does not accurately recapitulate modern human fat consumption, as lard availability has plummeted by nearly 80% over the past century, whereas vegetable oils – comprised of diverse fatty acids – are now the most common source of dietary fats in the United States^16^.

Over the last decade, it has become increasingly clear that consuming particular dietary fatty acids, rather than total fat, governs health and disease^17^. For example, both saturated fatty acids (SFAs) and trans fats are associated with worse cardiovascular outcomes, whereas monounsaturated (MUFAs) and polyunsaturated fatty acids (PUFAs) – such as those found in the Mediterranean diet – are beneficial to heart health^18–20^. These studies and resultant consensus guidelines^21^ have led to the cultivation and genetic modification of safflower, sunflower, soybean, and canola oils with high levels of the MUFA oleic acid (OA), and these high-oleic (HO) formulations are now widely available. Unlike heart disease, epidemiologic studies of HO fat intake have demonstrated variable risk reduction across cancer types^20,22^. Furthermore, efforts to link specific dietary fatty acids to PDAC risk have been hindered by recall bias, lack of granularity in diet surveys, or small sample size^14^. Together, these data argue that the role of dietary fatty acids in disease is context-specific such that there is not a “one size fits all” approach to disease prevention by diet modification.

Consistent with correlative findings in nutritional epidemiology research, preclinical studies have shown divergent effects of fatty acids on cancer progression. For example, OA drives melanoma lymph node metastasis^23^, the SFA palmitate (but *<u>not</u>* OA nor the PUFA linoleic acid (LA)) promotes oral carcinoma metastasis^24^, and LA enhances breast tumor growth^25^. In advanced PDAC models, diet modulation (*e.g.* calorie restriction vs. ketogenic diet) to reduce the MUFA/SFA ratio decreases tumor growth^26^. Most of these studies employed transplant models of cells derived from advanced cancers to elucidate how particular fatty acids govern the growth, maintenance, or metastasis of established tumors^27^. Therefore, whether and how specific dietary fats regulate tumor initiation and early cancer development is poorly understood.

To directly assess how dietary fat source and fatty acid composition influence PDAC development, we performed a comprehensive unbiased diet screen in an autochthonous oncogenic Kras-driven murine model using isocaloric HFDs differing solely in fat source and reflecting the breadth of modern human fat consumption^28^. We found that diets rich in the MUFA OA significantly enhanced pancreatic tumorigenesis, whereas diets enriched in the PUFA LA suppressed it. The quantity of dietary MUFAs and PUFAs governed the relative abundance of these fatty acids in pancreatic membrane phospholipids and regulated lipid peroxidation and sensitivity to ferroptosis. Collectively, our results demonstrate that dietary fat intake reshapes the pancreatic lipid landscape – specifically the balance of phospholipid saturation – to dictate pancreatic tumor development, suggesting that precise dietary factors may be able to alter cancer risk.

## RESULTS

### High fat consumption correlates with PDAC risk worldwide

Increasing PDAC incidence both in the United States and worldwide mirrors the overall increase in fat consumption over time^12^. To dissect the nuances of macronutrient contributions to PDAC, we examined correlations between diet availability documented by the Food and Agriculture Organization of the World Health Organization (WHO)^29^ and PDAC incidence rates from the Global Cancer Observatory (GLOBOCAN)^30^ across countries (**Table S1**). Dietary fat – as a percent of total calories and total fat availability – displayed the strongest association with increased PDAC incidence (**Figures 1A and 1B**). In contrast, carbohydrate availability, as a percentage of total calories, exhibited an inverse relationship with PDAC incidence, although the correlation disappeared when total carbohydrates were considered (**Figures 1C and 1D**). Protein availability showed a similar, although weaker, trend as fat availability (**Figures 1E and 1F**), likely due to a strong association between protein and fat availability globally (**Figure 1G**). However, fat accounts for a much greater quantity of calories than proteins across all populations. Accompanying the overall increase in fat consumption, there has been a shift in fat sources consumed such that vegetable oil-derived fat has become an increasingly large component of modern diets^12^. This has led to alterations in the proportion of different types of available and consumed dietary fatty acids, such that MUFAs (especially OA) have overtaken SFAs in the United States (**Figures 1H-1J**). Collectively, these correlations – based on worldwide population-level data – provide support for the hypothesis that HFD consumption, and potentially dietary fatty acid composition, may contribute to an altered risk of PDAC development.

**Figure 1:**
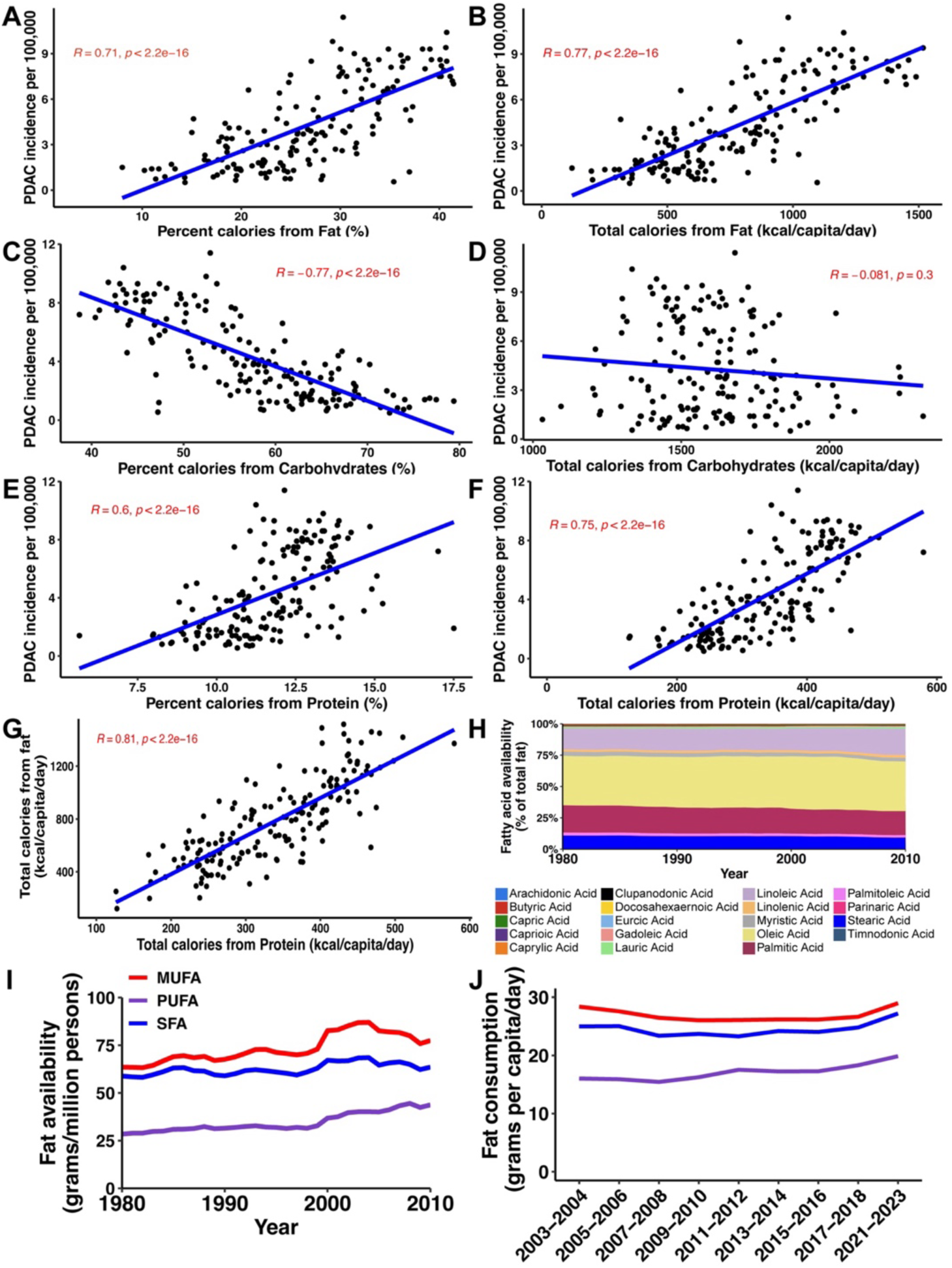
Fat availability correlates with increased PDAC incidence worldwide. (A) Positive correlation between % kcal fat availability (Food and Agriculture Organization (FAO) Food Balances) and age-standardized PDAC incidence (GLOBOCAN). Each point represents an individual country (n = 169). Spearman correlation coefficient (R) and *p*-value are shown. (B) Positive correlation between total calories from fat (kcal/capita/day) and PDAC incidence worldwide. (C) Negative correlation between the percentage of total calories from carbohydrates and PDAC incidence worldwide. (D) No correlation between total calories from carbohydrates (kcal/capita/day) and PDAC incidence worldwide. (E) Positive correlation between the percentage of total calories derived from protein and PDAC incidence worldwide. (F) Positive correlation between total calories from protein (kcal/capita/day) and PDAC incidence worldwide. (G) Positive correlation between total calories from fat (kcal/capita/day) and total calories from protein (kcal/capita/day). (H) Longitudinal analysis (1980-2010) of dietary fatty acid availability in the United States (United States Department of Agriculture (USDA)) shows that the MUFA oleic acid is the most abundant fatty acid. (I) Fat availability (grams per million persons) in the United States (sourced from the USDA) from 1980-2010 broken down by monounsaturated fatty acids (MUFA), polyunsaturated fatty acids (PUFA), and saturated fatty acids (SFA). (J) Fat consumption (grams per capita per day) in the United States (sourced from diet surveys conducted by National Health and Nutrition Examination Survey (NHANES)) from 2003-2023 broken down by MUFA, PUFA, and SFA.

### High oleic acid consumption enhances pancreatic tumorigenesis in mice

To systematically evaluate the effects of dietary fatty acid composition on pancreatic tumorigenesis, we leveraged an isocaloric HFD panel^28^, composed of twelve of the most commonly consumed fat sources in the United States (**Table S2**). These HFDs – modeled on the widely used 45% kcal lard HFD from Research Diets (D12451) – were identical in nutrient density, except for fat source, and better reflect patterns in modern human fat consumption (**Figures 1H-1J**). Importantly, the fat percentage approximated the upper end of average fat availability^13,15^ in countries in which PDAC incidence is the highest (**Figure 1A**). Furthermore, lipidomics analysis revealed that HFDs contained varying concentrations of saturated and unsaturated fatty acids (**Figure S1A**), which led to concordant changes in plasma fatty acid levels in HFD-fed wild-type (WT) C57BL/6 mice (**Table S3**), enabling us to correlate dietary and plasma fatty acid levels to pancreatic disease burden in PDAC models.

We fed this HFD panel to congenic C57BL/6 *Pdx1-Cre; Kras^LSL-G12D/+^* (KC) mice (**Figure S1B**), an oncogenic Kras-driven model that recapitulates the genetic and histologic progression of human PDAC from acinar-to-ductal metaplasia (ADM) to precursor pancreatic intraepithelial neoplasia (PanINs) to PDAC^31–33^. KC mice consistently gained weight over the 12-week feeding period (with the exception of mice fed fish oil, as has been previously described^34^) and consumed more HFD compared to a control diet (CD; 18% kcal soybean oil) but displayed only modest preferences between HFDs (average HFD intake of 12.9 kcal/mouse/day, range 11.3-14.6) (**Figures S1C and S1D**). Assessment of pancreatic disease burden (composite of ADM, PanINs, and PDAC) at the endpoint revealed that dietary fat source dictated tumorigenesis. Surprisingly, olive oil, HO safflower, and HO sunflower – HFDs rich in OA (**Figure S1A**) – significantly increased disease development in male KC mice (**Figure 2A**). Across the entire diet series, both dietary (**Figure S1A**) and plasma (**Table S3**) OA positively correlated with disease burden (**Figures 2B and 2C**). Interestingly, female mice exhibited less divergence in disease across diets with none of the HFDs significantly increasing tumor formation compared to CD (**Figure S2A**). This finding is concordant with reduced weight gain in females (**Figures S1D and** S**2B**) despite comparable food intake to males (average HFD intake of 11.9 kcal/mouse/day, range 10.9-13.4) (**Figures S1C and** S**2C**), as has been observed in prior HFD studies^35^. Furthermore, female mice did not show a correlation between dietary OA and disease burden (**Figure S2D**), arguing for sex-specific differences in the effect of OA on pancreatic tumor development.

**Figure 2:**
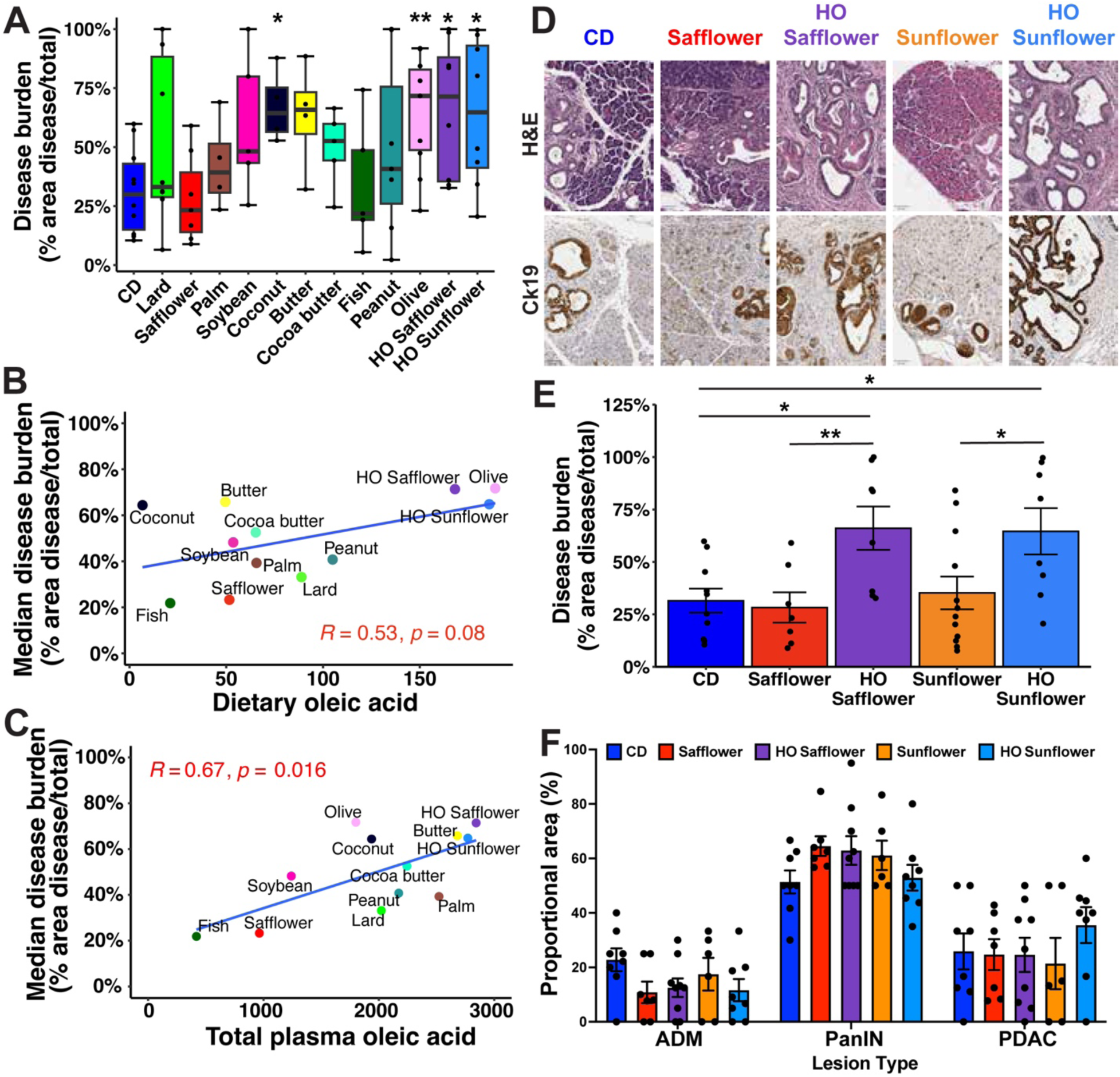
Oleic acid consumption promotes pancreatic tumorigenesis. (A) Disease burden (% disease/total pancreas area) for male KC mice fed control diet (CD) or designated HFDs for 12 weeks. Box plots designate 25^th^, 50^th^, and 75^th^ percentiles +/- min/max, n = 4-12 mice/diet. * p < 0.05, ** p < 0.01, Kruskal-Wallis test with Dunn’s post-hoc test. (B) Positive correlation between median disease burden of mice in (A) and dietary oleic acid concentration (µg/mg diet). Pearson correlation coefficient (R) and *p*-value are shown. (C) Stronger positive correlation between median disease burden of mice in (A) and mean total plasma oleic acid levels (nmol of FA/mL plasma) in non-tumor-bearing C57BL/6 male mice fed each diet for 12 weeks (n = 5 mice/diet). Pearson correlation coefficient (R) and *p*-value are shown. (D) Representative H&E and Ck19 (duct tumor marker) immunostaining of pancreata of male KC mice fed CD, LODs, or HODs for 12 weeks. Scale bars are 50 µm. (E) Disease burden (mean +/- s.e.m., n = 7-12 mice/diet) for male KC mice fed CD, LODs, or HODs for 12 weeks. * p < 0.05, ** p < 0.01, Kruskal-Wallis test with Dunn’s post-hoc test. (F) Proportional area (mean percentage +/- s.e.m., n = 6-9 mice/diet) of total disease by lesion type (ADM, PanIN, PDAC) of KC mice fed CD, LODs, or HODs. Proportions are not significantly different across diets, Kruskal-Wallis test with Dunn’s post-hoc test.

Due to the putative cardiovascular health benefits of OA, HO fats are readily available. This enabled us to directly evaluate the impact of OA on pancreatic tumorigenesis, by comparing low-oleic diets (LODs: safflower and sunflower) to cognate high-oleic diets (HODs: HO safflower and HO sunflower). Importantly, linoleic acid (LA), the predominant fatty acid in LODs, did not show a correlation with disease burden in KC mice (**Figure S1E**), providing a comparative baseline for the effects of OA. Strikingly, male (but not female) mice fed HODs exhibited a significant increase in disease burden compared to mice fed LODs (**Figures 2D, 2E, and S2E**). PanIN was the most frequent disease lesion across all diets (**Figure 2F**), arguing that HODs primarily increased the generation, expansion, and/or maintenance of PanINs, rather than promoting progression to more advanced disease. Consistent with this, the relative proportions of ADM, PanIN, and PDAC did not significantly differ by diet (**Figure 2F**), nor was metastatic disease evident in any mouse by the experimental endpoint. Although there was an overall positive correlation of disease burden with body weight across the HFD screen (**Figure S1F**), HO sunflower-fed male mice exhibited significantly increased disease burden (**Figures 2D and 2E**), despite no difference in body weight compared to sunflower-fed mice (**Figure S1D**). Finally, even though HO safflower-fed mice gained more weight than safflower-fed mice, body weight did not correlate with disease burden for either diet (**Figure S1G**). Collectively, these data argue that weight *per se* is not the primary driver of tumorigenesis and nominate dietary OA as potential contributor to PDAC development.

### Dietary fatty acids remodel pancreatic lipid metabolism and phospholipid composition

Multiple mechanisms have been implicated in obesity-driven tumorigenesis including (1) adipose-derived factors^36^, (2) pancreatic hormone dysregulation (hyperinsulinemia, β cell cholecystokinin (CCK) expression)^37,38^, increased tumor cell proliferation^39^, (4) induction of local inflammation and impaired T cell-mediated immunity^7,8,36,40^, and (5) modulation of cellular metabolism^7,10,26,41^. As WT C57BL/6 mice fed safflower and HO safflower HFDs for 12 weeks exhibited no significant differences in fat mass, fasting glucose, nor fasting insulin (**Figure S3**), diet-induced alterations in adiposity and glucose homeostasis likely did not significantly contribute to the pro-tumorigenic effects of OA. To further delineate the cellular mechanisms by which OA promotes PDAC progression, we analyzed pancreata from KC mice fed a control diet (CD, 18% kcal fat), LOD (safflower), or HOD (HO safflower). Since HODs principally augment Kras-driven PanIN development (**Figures 2D-2F**), we chose an early timepoint (3 weeks feeding when disease burden is comparable between dietary conditions (**Figure S4A**)) to allow a controlled comparison of low-grade PanIN lesions (**Figure S4B**) without the confounding effects of differences in tumor grade on tumor cell and microenvironmental features. Immunohistochemistry (IHC) revealed no significant differences in the proportion of Ki67+ and p19^ARF^+ PanIN cells comparing LOD vs. HOD (**Figure S4C**), arguing that OA does not modulate tumor cell proliferation nor senescence, respectively. As expected, HFD (vs. CD) increased F4/80+ macrophage-predominant immune cell infiltration (Cd45+) and smooth muscle actin-positive (SMA+) cancer-associated myofibroblasts, but this occurred to a similar extent in both HFD conditions (**Figures S4D and S4E**). Lymphocyte populations (B220+ B cells, Cd3+ T cells) were sparse and unchanged across diet conditions (**Figure S4F**). Together, these data argue that while HFDs had predictable effects on the fibroinflammatory microenvironment, these did not differ based on dietary fatty acid composition and thus are unlikely to contribute to the divergent effects of LODs and HODs on pancreatic tumorigenesis.

To determine unique effects of OA on pancreatic tumor biology, we performed bulk RNA-seq on pancreata from KC mice fed LODs (safflower, sunflower) vs. HODs (HO safflower, HO sunflower). Differential gene expression analysis (**Table S4**) revealed increased expression of genes involved in *de novo* lipogenesis (DNL), fatty acid elongation and desaturation, and membrane lipid remodeling in pancreata of mice fed HODs (**Figure S5A**). Gene set enrichment analysis (GSEA) confirmed significant enrichment of pathways associated with lipid biosynthesis and phospholipid remodeling (**Figure 3A**), suggesting a coordinated upregulation of anabolic lipid pathways in response to increased OA consumption. To determine whether these effects could be direct, we isolated primary acinar cells – the putative tumor cell-of-origin for PDAC^32,42^ – from WT C57BL/6 mice and treated them with exogenous fatty acids *in vitro*. OA (but not LA) rapidly (within 6 hours) induced the expression of key DNL genes (*Fasn* and *Scd2*) (**Figure S5B**), supporting the notion that OA stimulates anabolic lipid metabolism at the transcriptional level independent of oncogenic KRAS signaling or tumorigenic context. We functionally validated the ability of OA to induce DNL in murine *Kras^LSL-G12D/+^; Trp53^LSL-R172H/+^; Pdx1-Cre* (KPC) PDAC cells as OA treatment (in delipidated media) was sufficient to increase cellular levels of palmitate, the major lipid product of DNL (**Figure S5C**).

**Figure 3:**
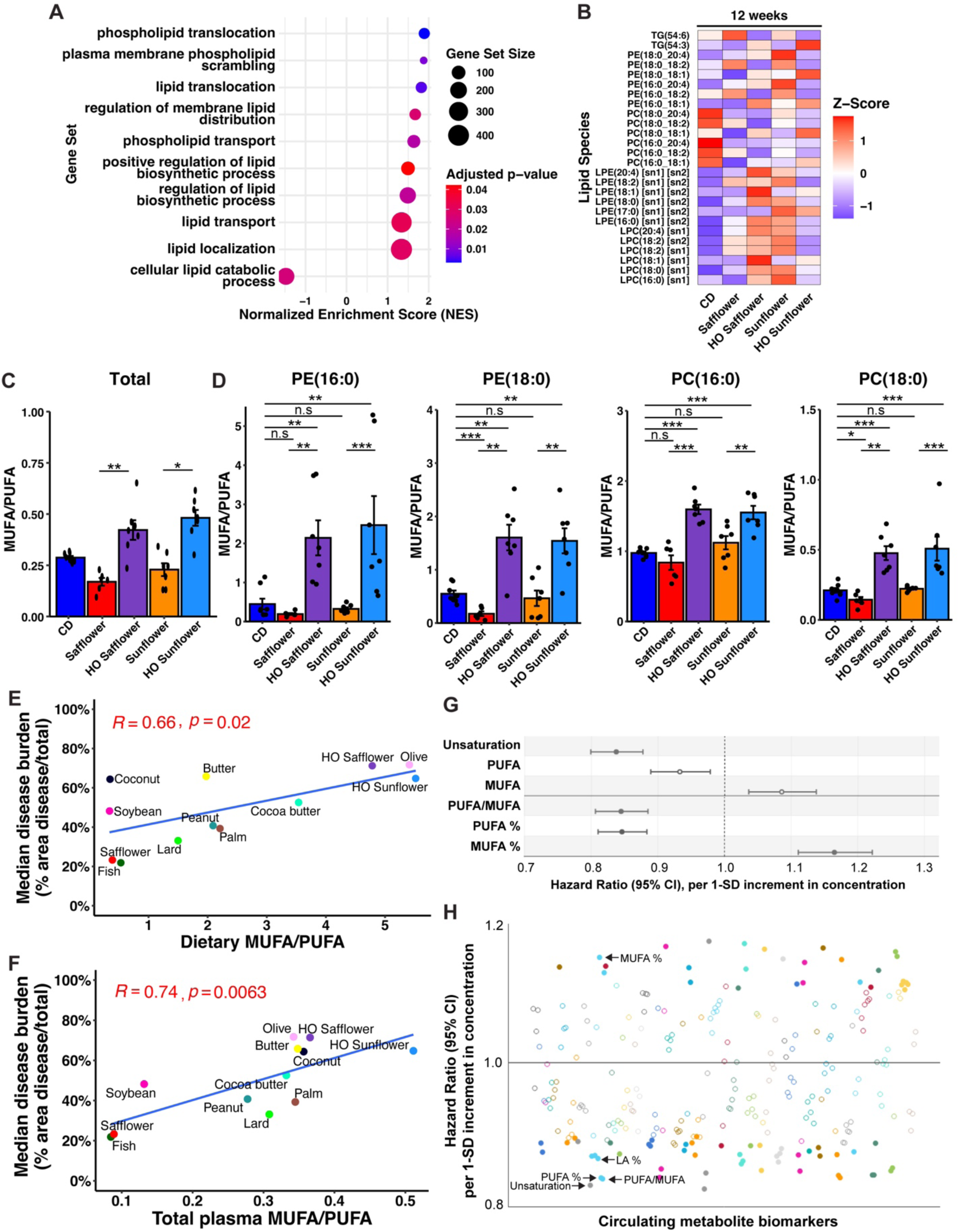
Dietary fatty acids remodel pancreatic lipid metabolism and phospholipid composition. (A) Gene set enrichment analysis (GSEA) of Gene Ontology Biological Process (GO:BP) gene sets related to lipid metabolism in the pancreata of KC mice fed HODs vs. LODs for 12 weeks (n = 8-10 mice/group). (B) Heatmap of lipid species detected by targeted LC-MS/MS-based lipidomics in the pancreata of male KC mice fed designated diets for 12 weeks (n = 6-8 mice/diet). Z-scores were generated using relative abundance, and rows are scaled. (C) Total MUFA/PUFA ratio (mean relative abundance +/- s.e.m. of OA(18:1)/LA(18:2) side chains across all PC, PE, LPC, LPE, and TG species measured) of pancreata of male KC mice in (B). * p < 0.05, ** p < 0.01, Kruskal-Wallis test with Dunn’s post-hoc test. (D) MUFA/PUFA ratio (mean relative abundance +/- s.e.m. of OA(18:1)/LA(18:2) side chains) in membrane phospholipids (PC and PE) of mice in (B). * p < 0.05, ** p < 0.01, Kruskal-Wallis test with Dunn’s post-hoc test. (E) Positive correlation between median disease burden in KC mice (n = 4-12 mice/diet) and dietary MUFA/PUFA (total esterified and non-esterified forms) across the diet series. Pearson correlation coefficient (R) and *p*-value are shown. (F) Strong positive correlation between median disease burden in KC mice and plasma MUFA/PUFA (total esterified and non-esterified forms) of non-tumor-bearing C57BL/6 male mice fed each diet for 12 weeks (n = 5 mice/diet). Pearson correlation coefficient (R) and *p*-value are shown. (G) Hazard ratio (HR) +/- 95% confidence interval (CI) for baseline circulating metabolites with PDAC incidence in the UK Biobank cohort (n = 487,689 with 1789 incident cases) adjusted for age, sex, and UK Biobank assessment center (Cox regression analysis). Statistically significant (*p* < 0.000005) HRs are denoted by filled in circles, including fatty acid unsaturation, PUFA/MUFA, PUFA%, and MUFA%. (H) Fatty acid unsaturation, PUFA/MUFA, PUFA%, and LA% were in the top 5% of measured metabolites (n = 249) in the UK Biobank cohort in (G) associated with the lowest HR for PDAC incidence, whereas MUFA% was in the top 5% of metabolites associated with the highest HR.

Given the transcriptional and functional effects of OA on lipid metabolism in pancreatic cells, we measured pancreatic lipids directly by targeted lipidomic analyses of HFD-fed KC mice (**Figure 3B; Table S5**). Major phospholipid classes – including phosphatidylcholine (PC) and phosphatidylethanolamine (PE), the most abundant phospholipids in mammalian tissues^43^ – exhibited significant shifts in relative abundance in response to HFD. In particular, we observed a ∼30% decrease in the PC/PE ratio in pancreata of HFD-fed KC mice (both LODs and HODs) compared to those on a control diet (CD) (**Figure S5D**). RNA-seq of *KC* pancreata revealed significantly increased expression of the rate-limiting enzyme for *de novo* PE synthesis (*Pcyt2*) in response to HFD (**Figure S5E**), which may contribute to this shift. In contrast, rate-limiting enzymes in *de novo* PC synthesis (*Pcyt1a*) and PE to PC conversion (*Pemt*) exhibited minimal changes between CD and HFD groups. Given prior reports linking low PC/PE ratios to increased lipid stress and metabolic dysregulation with HFD feeding^44–46^, these findings confirm that HFDs remodel tissue phospholipid homeostasis, which may contribute to their tumor-modulatory effects.

Due to marked divergence in tumor phenotypes (**Figures 2D and 2E**), we next focused on differences in pancreatic lipids between LOD- vs. HOD-fed KC mice, which primarily centered on the phospholipid fatty acid side chain abundance. Strikingly, the predominant fatty acids in the cognate diets (OA (18:1) in HODs vs. LA (18:2) in LODs) were enriched in PC and PE species, consistent with diet incorporation into pancreatic membrane phospholipids (**Figure 3B**). This led to significant changes in the ratio of lipids (PC, PE, LPC, LPE, and triglycerides (TG)) harboring MUFA (OA) and PUFA (LA) side chains (**Figures 3C and 3D)**. The differences were even more striking after just 3 weeks of feeding (**Figures S5F and S5G**), when disease burden was comparable across diets (**Figures S4A and S4B**). Specifically, LODs reduced – and HODs increased – the MUFA/PUFA side chain ratios in the most abundant membrane phospholipids (PC and PE) compared to CD (**Figures 3D and S5H**). We hypothesized that these diet-induced shifts in phospholipids containing MUFAs vs. PUFAs may contribute to tumorigenesis. Consistent with this hypothesis, both dietary and plasma MUFA/PUFA exhibited a stronger positive correlation with disease burden (**Figures 3E and 3F**) than OA, MUFA, PUFA, SFA, or MUFA/SFA (the latter of which was previously implicated in advanced PDAC tumor growth^26^) (**Figures 2B, 2C, and S5I**) across our diet series. As plasma fatty acid levels may be influenced by diet and correlate with the relative abundance of fatty acids in tissue phospholipids^47^, we analyzed the association between circulating fatty acids and incident PDAC risk in the UK Biobank^48^, the largest available prospective human cohort (487,689 individuals with 1789 incident PDAC cases) with rigorous plasma metabolite measurements. Multivariate Cox regression analyses revealed a significant association between baseline circulating MUFA% and increased risk of PDAC incidence (**Figure 3G**). Conversely, baseline circulating PUFA/MUFA, PUFA%, and overall fatty acid unsaturation were associated with a significantly decreased risk of incident PDAC (**Figure 3G**). Across all 249 circulating metabolites measured in the cohort, these fatty acid measures were in the top 5% of biomarkers associated with increased or decreased PDAC risk (**Figure 3H**). Collectively, these data support the notion that diet-induced imbalances in phospholipid fatty acid saturation – specifically MUFAs and PUFAs – may drive PDAC development.

### Opposing effects of exogenous MUFAs and PUFAs on lipid peroxidation and ferroptosis sensitivity in pancreatic tumor cells in vitro

Lipids are critical mediators of membrane fluidity, bioenergetics, and signaling, supporting the importance of lipid composition – particularly the level of saturation – in establishing and maintaining the cancer cell phenotype^49,50^. For example, excessive oxidation of PUFA-containing phospholipids (in particular PE linked to arachidonic acid (20:4) or adrenic acid (22:4)^51^, the elongation products of LA (18:2)), results in ferroptosis, characterized by the iron-dependent accumulation of lipid peroxides and oxidative damage to cell membranes^52–55^. Unlike PUFAs, MUFAs are resistant to oxidation, and therefore, cells with MUFA-rich membranes are less likely to undergo ferroptosis. Consistent with this, endogenous OA in lymph protects melanoma metastases from ferroptosis^23^, and exogenous OA confers ferroptosis resistance in cultured cells, worms, and mouse liver^56,57^. We validated this model in human *KRAS^G12D^* mutant KP4 PDAC cells treated with OA alone or in combination with equimolar LA in the presence or absence of GPX4 inhibitors (RSL3 and ML162), which induce ferroptosis. In basal conditions, GPX4 inhibitor-treated KP4 cells exhibited a dose-dependent increase in lipid peroxidation (**Figure 4A**). OA treatment significantly rescued KP4 cells from loss of cell viability induced by GPX4 inhibitors, but this protective effect was abolished by the addition of LA (**Figure 4B**).

**Figure 4:**
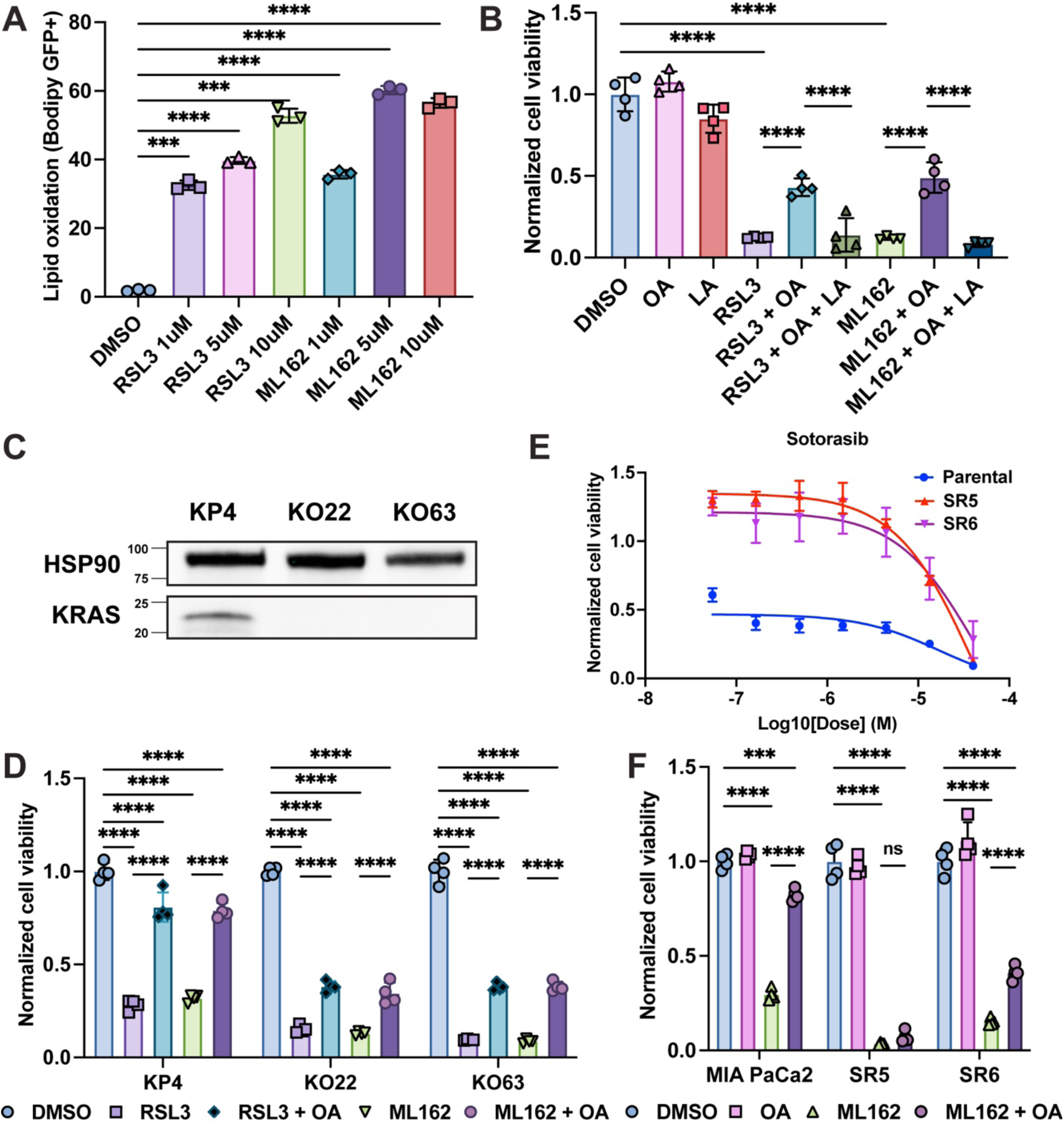
OA induces ferroptosis resistance in PDAC cells in a mutant KRAS-dependent manner. (A) Quantification of lipid peroxidation (C11-BODIPY(581/591) oxidized fluorescence at 488 nm, mean +/- s.d., n = 3 technical replicates) in human PDAC KP4 cells treated with increasing doses of the GPX4 inhibitors (RSL3, ML162) vs. vehicle (DMSO) for 3 hours. ***p < 0.001, and ****p < 0.0001, one-way ANOVA with Tukey’s post-hoc test. (B) Cell viability (normalized to DMSO, mean +/- s.d., n = 4 technical replicates) of KP4 cells treated with RSL3 (0.1µM) or ML162 (0.073µM) with or without exogenous fatty acids oleic acid (OA, 200 µM) and linoleic acid (LA, 200 µM). ****p < 0.0001, one-way ANOVA with Tukey’s post-hoc test. (C) Western blot confirms loss of KRAS protein in CRISPR-mediated knockout clones (KO22, KO63) compared to parental KP4 cells. HSP90 is a loading control. (D) Cell viability (normalized to DMSO, mean +/- s.d., n = 4 technical replicates) of KP4, KO22, and KO63 cells treated with RSL3 (0.1µM) or ML162 (0.073µM) with or without OA (200 µM). **** p < 0.0001, one-way ANOVA with Tukey’s post-hoc test. (E) Dose-response curves (normalized cell viability to DMSO, mean +/- s.d., n = 4 technical replicates) for KRAS^G12C^ inhibitor sotorasib of MiaPaCa-2 parental cells and two sotorasib-resistant clones (SR5 and SR6). (F) Cell viability (normalized to DMSO, mean +/- s.d., n = 4 technical replicates) of MiaPaCa-2, SR5, or SR6 cells treated with ML162 (0.073µM) with or without OA (200 µM). SR5 and SR6 were maintained on continuous sotorasib (1 µM) treatment. *** p<0.001, **** p < 0.0001, one-way ANOVA with Tukey’s post-hoc test. n.s. = non-significant.

Prior studies have reported that mutant RAS enhances fatty acid scavenging^58,59^, and *KRAS* mutant pancreatic cells may have enhanced capacity to modulate lipid peroxidation and ferroptosis pathways via uptake of exogenous fatty acids. To determine whether mutant *KRAS* is required for OA-mediated ferroptosis resistance, we studied *KRAS*-deficient KP4 cells generated by CRISPR-Cas9-mediated knockout, as previously described^60^ (**Figure 4C**). *KRAS*-deficient clones exhibited greater sensitivity to ferroptosis inducers (**Figure 4D**), concordant with previous studies demonstrating increased vulnerability of therapy-resistant cells to GPX4 inhibitors^61^. Nonetheless, OA partially rescued cell viability in *KRAS*-deficient cells (**Figure 4D**), suggesting that OA-mediated ferroptosis resistance does not strictly depend on mutant KRAS. However, the absolute increase in cell viability was greater when mutant *KRAS* was retained, consistent with KRAS-dependent enhancement of the effects of exogenous OA (**Figure 4D**). As an orthologous approach, we tested whether pharmacologic KRAS inhibition could blunt the protective effects of exogenous OA on ferroptosis sensitivity. We generated *KRAS^G12C^* mutant MiaPaCa-2 clones that survived sustained mutant KRAS inhibition with the FDA-approved allele-specific KRAS^G12C^ inhibitor sotorasib (**Figure 4E**). Sanger sequencing of KRAS exons did not identify secondary *KRAS* mutations in resistant clones, which would bypass KRAS inhibition. Like *KRAS* knockout, mutant KRAS inhibition reduced the capacity of OA to sustain cell viability in the presence of GPX4 inhibition (**Figure 4F**). Collectively, these findings suggest that exogenous OA protects pancreatic tumor cells against ferroptosis, that this effect can be enhanced by mutant KRAS, and that LA can reverse this effect, implicating the balance of exogenous MUFAs and PUFAs in regulating ferroptosis sensitivity.

### Dietary fatty acids govern lipid oxidation and ferroptosis sensitivity in the pancreas

We next evaluated whether similar alterations in ferroptosis sensitivity could be induced by dietary fatty acids *in vivo*. Functional over-representation analysis of RNA-seq data revealed an enrichment of pathways associated with lipid oxidation in pancreata of mice fed LODs vs. HODs for 12 weeks (**Figure 5A**). Furthermore, GSEA revealed that as little as 3 weeks of LOD feeding caused significant enrichment of an established ferroptosis transcriptional signature (genes induced by erastin in HT-1080 cells^52^) (**Figures 5B and S6A**). These data suggest that PUFA-rich LODs (safflower, sunflower) may prime pancreatic phospholipids for lipid peroxidation and ferroptosis susceptibility, whereas MUFA-rich HODs (HO safflower, HO sunflower) may counter these effects. Consistent with this hypothesis, LODs increased the levels of 4-hydroxynonenal (4-HNE), a lipid peroxidation byproduct, and the oxidative stress marker Nqo1, a major enzyme in the cellular antioxidant response, in PanIN lesions compared to HODs and CD (**Figures 5C, 5D**, S**6B, and S6C**). Targeted lipidomics further revealed that mice fed LOD exhibited a prompt (within 3 weeks) enrichment of PE(20:4), a major mediator of ferroptosis^51^, compared to mice fed HOD (**Figure 5E**). These results indicated that LODs may enhance ferroptosis sensitivity in the pancreas *in vivo*. Given the lack of specific and reliable pharmacologic agents for *in vivo* ferroptosis induction, we instead fed KC mice CD, LOD (safflower HFD), or HOD (HO safflower HFD) for 3 weeks, isolated primary acinar cells, and tested ferroptosis sensitivity *ex vivo*. As predicted, LOD feeding increased sensitivity to GPX4 inhibitors relative to CD and HOD (**Figure 5F)**, showing directly that dietary LODs enhance ferroptosis sensitivity in the pancreas. These data argue that dietary fatty acids remodel pancreatic cell membrane phospholipids to perturb lipid peroxidation and oxidative stress pathways that govern ferroptosis susceptibility.

**Figure 5:**
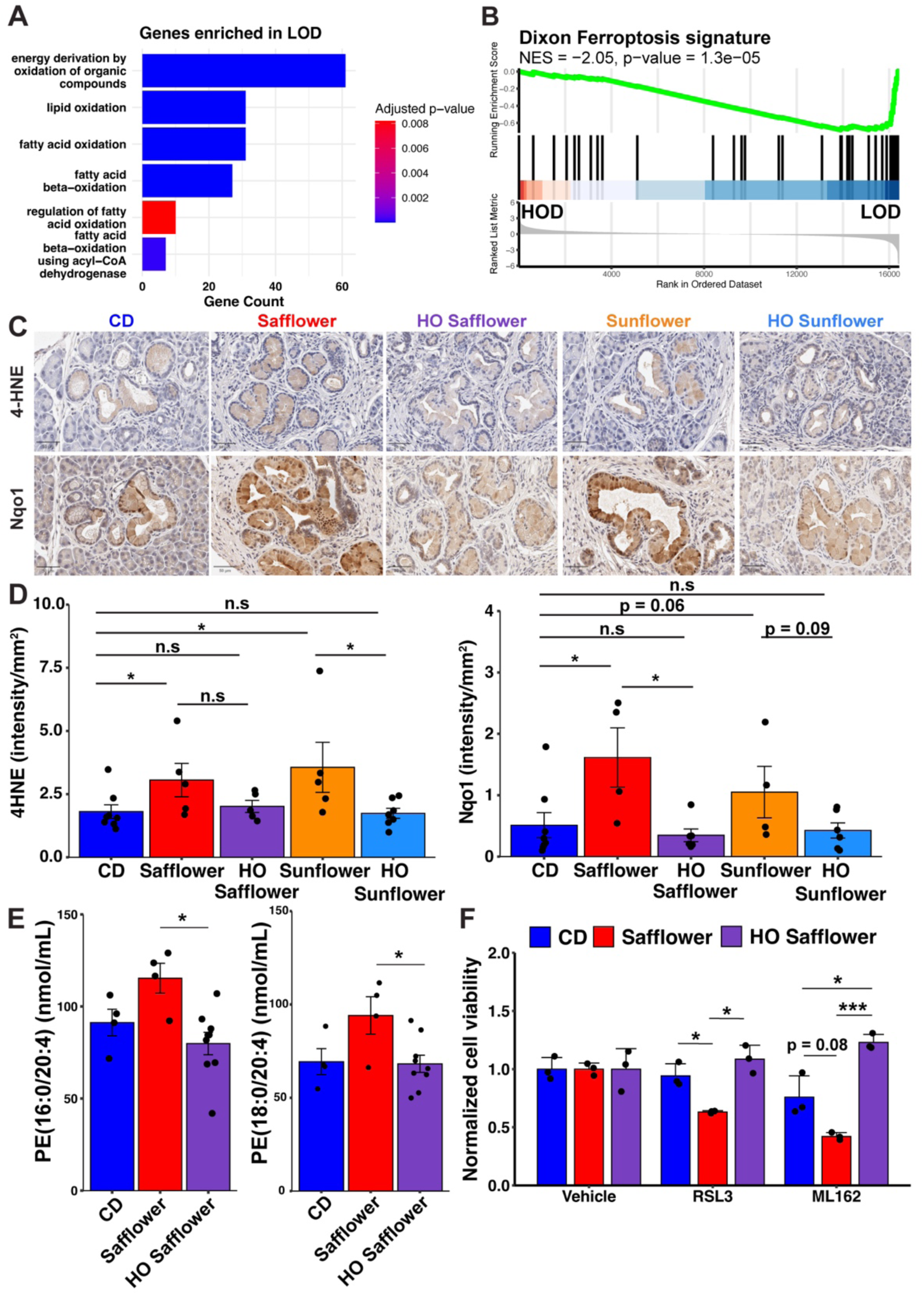
Dietary fatty acid composition governs pancreatic lipid oxidation and ferroptosis sensitivity. (A) Functional overrepresentation analysis shows enrichment of transcriptional signatures of lipid oxidation (Gene Ontology (GO) Biological Processes) in the pancreata of KC mice fed LODs vs. HODs for 12 weeks (n = 8-10 mice/group). (B) GSEA reveals enrichment of the Dixon Ferroptosis signature (erastin-treated HT-1080 cells^52^) in pancreata of KC mice in (B) with LOD feeding. Normalized enrichment score (NES) and *p*-value are shown. (C) Representative images of immunohistochemical staining for 4-hydroxynonenal (4-HNE) (top row) and Nqo1 (bottom row) of PanINs in KC mice fed the designated diets for 12 weeks. Scale bars are 50 µm. (D) Quantification of IHC in (C) denoting intensity/mm^2^ (mean +/- s.e.m.) in PanIN lesions for each immunostain. * p < 0.05, Kruskal-Wallis test with Dunn’s post-hoc test. (E) Concentrations (mean +/- s.e.m.) of phosphatidylethanolamine (PE) lipid species containing arachidonic acid side chains (16:0/20:4 and 18:0/20:4) in the pancreata of KC mice fed designated diets (n = 4-9 mice per diet). * p < 0.05, one-way ANOVA with Tukey’s post-hoc test. (F) Normalized cell viability (mean +/- s.d., normalized to vehicle) of primary acinar cells harvested from male KC mice fed CD, safflower HFD, or HO safflower HFD for 3 weeks (n = 2 pooled mice/diet) and treated with GPX4 inhibitors (RSL3 or ML162) vs. vehicle (DMSO) for 6 hours. Dots represent technical replicates (n = 3) from a representative biologic replicate. * p < 0.05, *** p < 0.001, Kruskal-Wallis test with Dunn’s post-hoc test.

### Increased dietary polyunsaturated fatty acid intake suppresses PDAC development

Since ferroptosis is a potential tumor suppressive mechanism^62,63^, we hypothesized that increased PUFA consumption – by providing substrates for lipid peroxidation and ferroptosis *in vivo* – would impede PDAC progression. In KC mice, high PUFA HFDs (LODs) showed a non-significant trend towards decreased PanIN formation compared to CD (**Figure 2D and 2E**) despite increased fat intake and weight gain (**Figure S1C and S1D**). We reasoned that the tumor suppressive effects of increased dietary PUFA intake may have been masked due to the low overall disease burden in KC mice decreasing the ability to detect a significant reduction. Furthermore, endogenous MUFA production would be expected to counter the effects of dietary PUFAs, unless consumed at high levels. To circumvent this, we leveraged our previously reported KCO (*Kras^LSL-G12D/+^; Pdx1-Cre; Lep^ob/ob^*) model, characterized by rapid onset obesity and markedly accelerated PDAC progression due to overating^37^. The propensity of this mouse model to voraciously consume calories provided a powerful approach to enrich the pancreas with PUFAs from the diet. KCO mice fed a safflower HFD (PUFA-rich LOD) for 8 weeks exhibited significantly increased weight gain and caloric intake relative to CD-fed mice (**Figures 6A, 6B, and S6D**). As expected, lipidomics analyses (**Table S5**) confirmed a significant decrease in the MUFA/PUFA ratio in membrane phospholipids, concordant with increased 4-HNE and transcriptional signatures of lipid oxidation (**Figures 6C and S6E-S6H**). Strikingly, safflower HFD-fed KCO mice exhibited significantly reduced disease burden, with the majority of mice harboring neoplastic lesions stalled in low-grade PanIN stages (**Figures 6D and 6E**). Importantly, disease burden was reduced in males and females, concordant with the capacity of safflower HFD feeding to enhance *ex vivo* ferroptosis sensitivity in primary acinar cell cultures derived from mice of both sexes (**Figures 5F**, **S2F, and S2G**). Overall, there was an inverse correlation between weight gain – typically associated with increased PDAC risk in humans and murine models – and disease burden (**Figure 6F**). Interestingly, safflower HFD-fed KCO mice exhibited greater variance in overall disease burden (**Figure 6D**). We suspected that differences in fatty acid consumption and/or incorporation in the pancreas between mice could account for this variation. Consistent with this hypothesis, disease burden in KCO mice was inversely correlated with 18:2 (LA) containing phospholipids in the pancreas (**Figure 6G**), as measured by lipidomics analysis. Furthermore, the total MUFA/PUFA ratio of phospholipids was strongly correlated with tumorigenesis (**Figure 6H**). Finally, pancreatic RNA-seq analysis demonstrated that safflower HFD-fed KCO mice with lower disease burden exhibited significant enrichment in transcriptional signatures of ferroptosis compared to those with higher disease burden (**Figure 6I**). These results argue that dietary PUFAs act as suppressors of PDAC progression by increasing availability of PUFA-containing phospholipids for peroxidation and ferroptosis.

**Figure 6:**
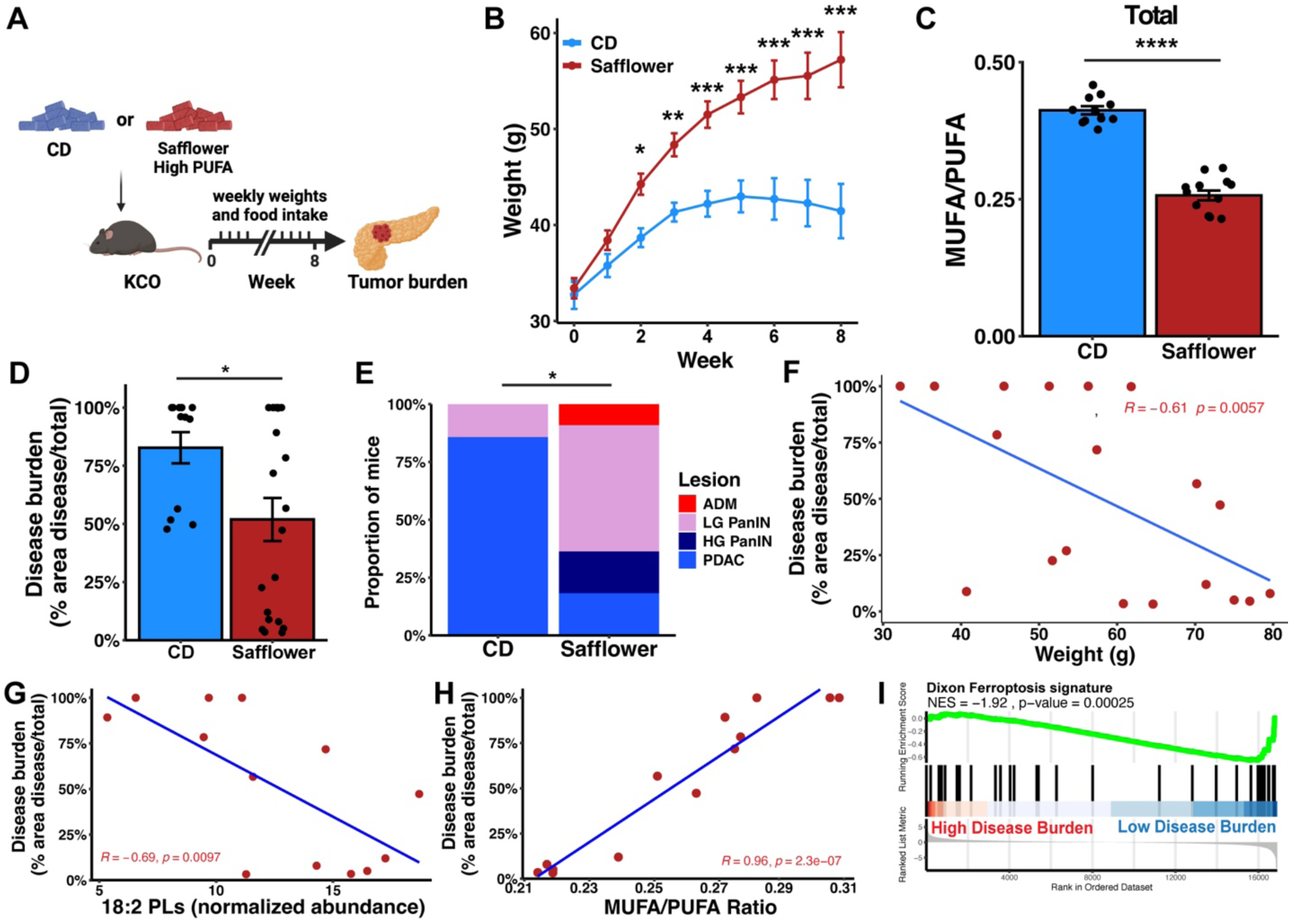
Increased dietary polyunsaturated fatty acid intake suppresses PDAC development. (A) Experimental schema for KCO feeding experiment. KCO mice were randomized to control diet (CD) or PUFA-rich LOD (safflower HFD) at 6 weeks of age. After 8 weeks HFD feeding, mice were analyzed for tumor phenotypes. (B) Body weight (grams (g), mean +/- s.e.m.) of KCO mice fed CD or safflower HFD over 8 weeks (n = 15-27 mice/diet). * p < 0.05, ** p < 0.01, *** p < 0.001, repeated measures ANOVA (rmANOVA) followed by post-hoc comparisons using estimated marginal means (EMMeans). (C) Total MUFA/PUFA ratio (mean relative abundance +/- s.e.m. of OA(18:1)/LA(18:2) side chains across all PC, PE, LPC, LPE, and TG species measured) of pancreata of KCO mice (n = 11-13 mice/diet) fed CD or safflower HFD for 8 weeks. **** p < 0.001, Wilcoxon rank-sum test. (D) Disease burden (mean +/- s.e.m.) for KCO mice fed CD or safflower HFD for 8 weeks (n = 12-20 mice/diet). * p < 0.05, Wilcoxon rank-sum test. (E) Proportion of mice with the highest-grade lesion categorized as acinar-to-ductal metaplasia (ADM), low-grade pancreatic intraepithelial neoplasia (LG PanIN), high-grade PanIN (HG PanIN), or PDAC for mice in (D). * p <0.05, Fisher’s exact test. (F) Negative correlation between body weight and disease burden for mice fed safflower HFD for 8 weeks. Pearson correlation coefficient (R) and *p*-value are shown. (G) Negative correlation between pancreatic LA(18:2)-containing phospholipids (PLs) and disease burden for KCO mice fed safflower HFD for 8 weeks. Pearson correlation coefficient (R) and *p*-value are shown. (H) Positive correlation between the total pancreatic lipid MUFA/PUFA ratio (18:1(OA)/18:2(LA) of all PC, PE, LPC, LPE, and TG species measured) with disease burden in KCO mice fed safflower HFD for 8 weeks. Pearson correlation coefficient (R) and *p*-value are shown. (I) GSEA reveals enrichment of the Dixon Ferroptosis Signature (erastin-treated HT-1080 cells^52^) in pancreata of KCO mice fed safflower HFD with low (<50%) vs. high (>50%) disease burden. Normalized enrichment score (NES) and *p*-value are shown.

### Loss of p53 overrides the impact of dietary fatty acids on PDAC development

Genetic loss of p53 function is a frequent event in PDAC progression, occurring in >50% of human PDAC tumors^64,65^. Through transcriptional regulation of cysteine transport, glutathione (GSH) production, or cell cycle state, p53 has been shown to either promote or repress ferroptosis in different contexts and in response to different ferroptotic stimuli (cysteine deprivation vs. GPX4 inhibition)^66–69^. Therefore, we sought to determine whether the tumor suppressive or promoting effects of LODs and HODs, respectively, could be modulated by loss of p53. To study how p53 loss alters the impact of HFD on PDAC progression, we took advantage of the K-MADM-p53 model we developed, which enables lineage tracing of genetically distinct subclonal populations of cells retaining or lacking p53 within tumors initiated by oncogenic Kras expression^70^. This model – based on the mosaic analysis with double markers (MADM) system^71,72^ – generates tandem dimer Tomato-expressing (tdTomato+) p53 wild-type (p53^WT^) cells and green fluorescence protein-expressing (GFP+) p53-deficient (p53^null^) cells that are born at a 1:1 ratio from a single Cre-mediated mitotic *inter*-chromosomal recombination event (**Figure 7A**). By counting the proportions of GFP+ and tdTomato+ low-grade PanINs, high-grade PanINs, and PDAC lesions, we previously found that p53 loss enhances PanIN initiation, proliferation, and progression to advanced disease^70^. Importantly, since the p53^WT^ and p53^null^ clones reside in the same mouse and thus are exposed to the same dietary influence, we can rigorously assess how p53 influences the impact of diet on tumorigenesis (**Figure 7B**).

**Figure 7:**
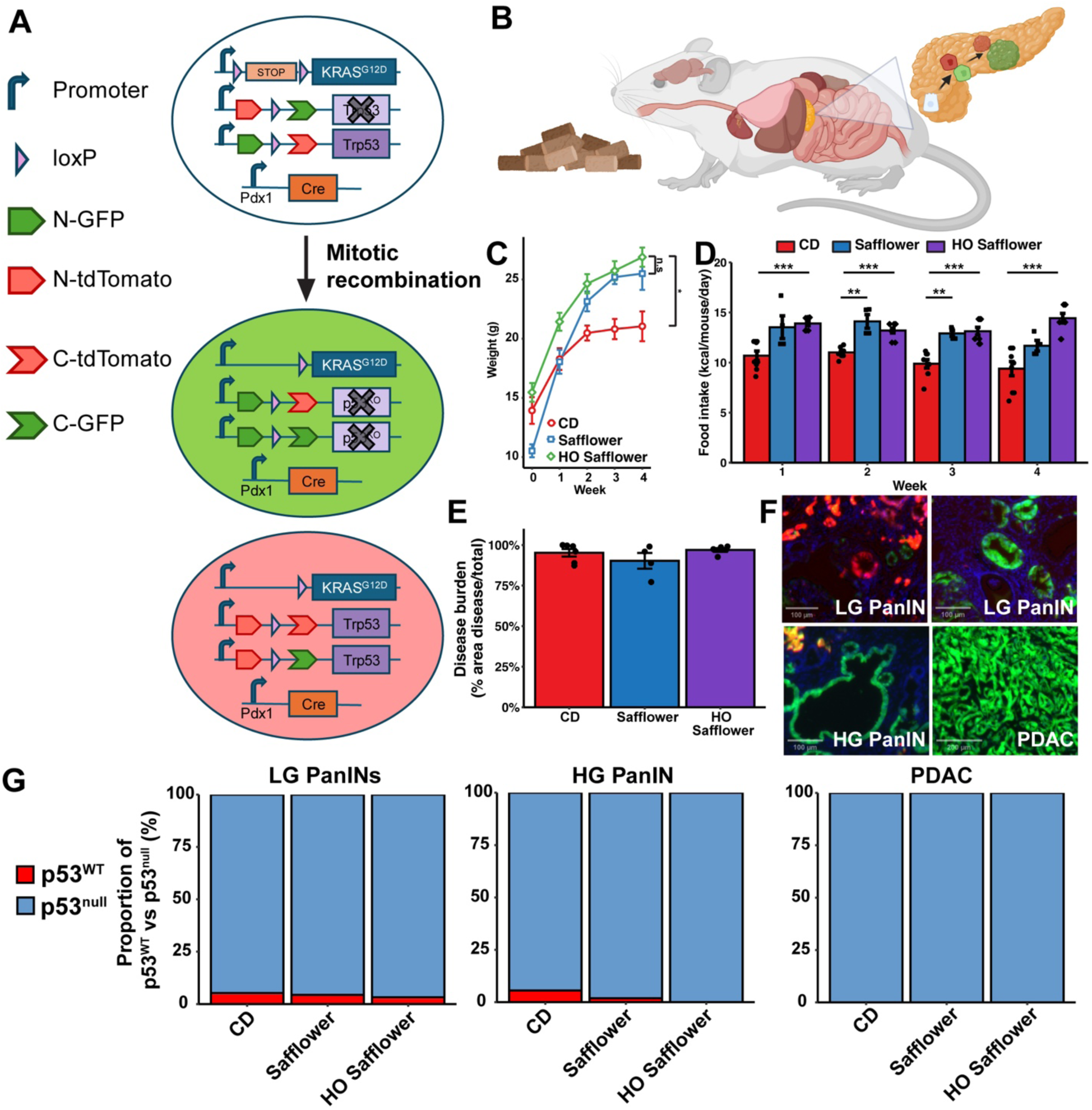
p53 loss overrides the impact of dietary fatty acids on PDAC development. (A) Schematic of MADM system that generates GFP+ p53^null^ and tdTomato+ p53^WT^ cells from a rare Cre-mediate mitotic *inter*-chromosomal recombination event in *Kras* mutant pancreatic epithelial cells^70^. (B) Tumor development of GFP+ p53^null^ and tdTomato+ p53^WT^ cells can be monitored within the same mouse subject to diet treatment. Created with BioRender.com. (C) Body weight (grams (g), mean +/- s.e.m.) of K-MADM-p53 mice fed 18% kcal control diet (CD) or 45% kcal HFDs (safflower or HO safflower) for 4 weeks (n = 4-9 mice/diet). * p < 0.05 for each HFD vs. CD, Kruskal-Wallis test with Dunns post-hoc test. (D) Food intake (kcal/mouse/day, mean +/- s.e.m.) for K-MADM-p53 mice in (C). ** p < 0.01, *** p < 0.001, Kruskal-Wallis test with Dunns post-hoc test. (E) Disease burden (mean +/- s.e.m.) of mice at endpoint for K-MADM-p53 mice in (C). (F) Representative low-grade PanINs (LG PanINs), high-grade PanINs (HG PanINs), and PDAC labelled with MADM fluorescent markers. Scale bars are 100 µm. (G) Proportion of tdTomato+ p53^WT^ (red) vs. GFP+ p53^null^ (blue) cells of K-MADM-p53 mice in (C), categorized as LG PanINs, HG PanINs, or PDAC by pathologic analysis. No statistically significant differences were observed between HFDs and CD for each lesion type (Fisher’s exact test).

We fed K-MADM-p53 mice CD, safflower HFD (PUFA-rich LOD), or HO safflower HFD (MUFA-rich HOD) for 4 weeks after weaning and analyzed fluorescence in the pancreas. Although mice fed HFDs displayed increased weight gain and food intake relative to CD-fed mice (**Figures 7C and 7D**), all mice developed significant disease burden irrespective of diet (**Figure 7E**). As expected^70^, CD-fed mice exhibited increased GFP+ (vs. tdTomato+) low-grade PanIN, high-grade PanIN, and PDAC lesions (**Figures 7F and 7G**). Strikingly, HFD feeding – regardless of MUFA or PUFA predominance – did not alter these tumor phenotypes (**Figure 7G**). These data argue that p53 loss overrides the effects of diet on tumor progression, consistent with the potential for a dominant impact of somatic genetic alterations over dietary influence in pancreatic tumorigenesis.

## DISCUSSION

In this study, we leveraged a large well-controlled HFD panel in a well-defined autochthonous PDAC model to identify a previously unrecognized role for dietary fatty acid composition in pancreatic tumorigenesis. Combined with our prior work demonstrating that overeating-induced obesity (*Lep^ob/ob^*) promotes PDAC progression^37^, we find that both diet *quantity* and diet *quality* mediate pancreatic tumor development. Unlike high BMI, which has uniformly been associated with increased PDAC risk in prospective human cohorts^73^, the reported associations between HFD consumption and PDAC have not been consistent^74^. Our results suggest that specific differences in fatty acid composition may be more important than overall HFD intake in modulating pancreatic tumorigenesis, leading to these mixed results. Epidemiologic studies in humans examining the association of specific fatty acid classes (SFA, MUFA, PUFA) with PDAC risk have also shown discordant findings, likely due to differences in study design and confounding variables, such as demographic and genetic characteristics of the study population, sample size, respondent and recall bias in dietary questionnaires, and limited follow-up^14^. Our findings in mice offer clarity on the complex impacts of diverse fatty acids on tumorigenesis and implicate the balance of fatty acid consumption (MUFA (OA) vs. PUFA (LA)) as a more significant contributor to PDAC development than any single fatty acid. Moreover, our work overcomes many of the important limitations of prior preclinical studies of diet and cancer^12–14^ and validates the power of rigorous, systematic diet studies in mice to uncover unexpected links between dietary components and cancer.

Our results also demonstrate the profound extent to which dietary fat composition reshapes the tissue lipidome, as the fatty acid profile of the diet dictated the phospholipid landscape of the pancreas. Given the pleiotropic effects of different fatty acids on cellular function – including membrane dynamics, signaling pathways, and cell fate^49,50^ – shifting dietary fatty acid composition emerges as a powerful strategy to modulate tumor initiation and progression. We found that increased consumption of the MUFA OA enhanced tumor development, while the PUFA LA inhibited it through alterations in lipid peroxidation and ferroptosis sensitivity in the pancreas. Our findings align with the observation that the cellular antioxidant response is critical for oncogenic *Kras*-driven pancreatic tumorigenesis^75,76^. Prior work in mice showed that dietary MUFAs reprogram lipid metabolism to promote the growth of advanced PDAC tumors^26^, whereas cysteine depletion-mediated ferroptosis induction suppresses it^62^. Our study directly links diet-induced tissue lipid remodeling to alterations in ferroptosis sensitivity, demonstrates that these phenotypes can also perturb tumor initiation and early progression, and defines how this knowledge can be leveraged to restrain pancreatic tumorigenesis through consumption of a PUFA-rich diet. These results may have implications for nutritional guidance to individuals at high risk of developing PDAC, such as those with a strong family history or genetic predisposition^6^. Given the rising prevalence of HO fats in modern diets, increased MUFA consumption may inadvertently shift phospholipid composition, priming the pancreas for tumorigenesis. Longitudinal analyses of the UK Biobank cohort using objective measurements of circulating metabolites supports this hypothesis. If additional prospective epidemiologic and interventional studies validate these findings in human populations, our results would argue that what may be good for the heart, may not be so for the pancreas.

Ferroptosis has emerged as a powerful tumor suppressive mechanism with several clinical trials currently testing non-specific ferroptosis-inducing agents in cancer^63,77^. Our findings reinforce this growing body of evidence while elevating its translational relevance by demonstrating that diet can modulate ferroptosis sensitivity *in vivo*. Prior work has suggested that exogenous MUFAs reduce ferroptosis sensitivity in advanced cancer cells *in vitro*^56^. Conversely, lipid limitation induces TG lipolysis and PUFA liberation to confer ferroptosis sensitivity^78^. These studies imply that too much or too little fatty acid availability affects ferroptosis susceptibility. We show that these observations also pertain to carcinogenesis *in vivo*, as the saturation state of dietary fatty acids (MUFAs vs. PUFAs) lead to concordant changes in the fatty acyl side groups of pancreatic membrane phospholipids to govern lipid peroxidation, ferroptosis sensitivity, and tumorigenesis in the pancreas. This may be particularly relevant to PDAC development, as our data and those of others support a role for mutant KRAS in fatty acid uptake and phospholipid remodeling^58,59^. Collectively, these results suggest that dietary interventions – such as increasing PUFA intake – could serve as practical, non-pharmacologic approaches for cancer prevention by enhancing ferroptosis susceptibility in emerging pancreatic neoplasia.

Finally, there are some notable limitations of our study. Due to the challenges in interpreting nutritional epidemiology studies, as described above, we chose to pursue a well-controlled HFD screen^28^ in a autochthonous mouse model^31^ to clarify the relationship between dietary fatty acids and PDAC development. As noted above, additional epidemiologic or intervention studies will be required to confirm the relevance of these findings to humans. We also observed sex-specific differences in OA-driven tumorigenesis, which remain unexplained. Our results align with previous studies demonstrating sex-dependent responses to HFD feeding^35^. Notably, DNL has been reported to be lower in females^79^, suggesting that reduced endogenous MUFA production (compared to males) may lead to insufficient overall tissue MUFA levels to suppress ferroptosis despite increased OA consumption. In contrast to MUFAs, PUFAs are largely acquired by the diet. As a result, our data showed that high PUFA consumption enhanced ferroptosis sensitivity and suppressed tumorigenesis in mice irrespective of sex. Alternatively, it is possible that the sex-specific differences in the effect of increased OA consumption on tumorigenesis may be abrogated at thermoneutrality, as has been observed previously with HFD-induced metabolic dysfunction-associated steatotic liver disease in mice^80^. Future studies should explore the mechanistic underpinnings of these sex-specific effects of fatty acid consumption in humans to refine potential dietary recommendations. Finally, our work focused on controlling dietary macronutrients (fats, carbohydrates, and protein), but we did not fully account for the spectrum of micronutrients (vitamins and minerals) that may differ based on fat source and even the commercial vendor from which fats and oils are sourced. Although these micronutrients could play modifying roles in tumorigenesis, the strong correlations between fatty acid consumption and phospholipid remodeling, lipid peroxidation, ferroptosis sensitivity, and tumor progression argue that dietary fatty acid unsaturation dictates pancreatic tumorigenesis.

## RESOURCE AVAILABILITY

### Lead contact

Further information and requests for reagents or resources should be directed to and will be fulfilled by the lead contact, Mandar Deepak Muzumdar.

### Materials availability

All unique/stable reagents generated in this study will be provided by the lead contact upon publication with a completed materials transfer agreement.

### Data and code availability

RNA-seq data have been deposited in the Gene Expression Omnibus (GEO) with accession number GSE292815 and will be publicly available as of the date of publication. Lipidomic datasets generated during and/or analyzed during the current study will be submitted to the Metabolomics Workbench in LIPID MAPS and will be publicly available as of the date of publication. No unique code was generated for this study.

## ACKNOWLEDGEMENTS

We thank the Muzumdar lab members for helpful discussions and feedback; D. DiMaio and E. Forrest for critical reading of the manuscript; the Yale Keck DNA Sequencing Core and MGH CCIB DNA Core for DNA sequencing; the Yale Center for Genome Analysis (YCGA) for library preparation and RNA-sequencing; and Drs. T. Jacks, A. Lowy, and L. Luo for mice. C.F.R. was supported by a postdoctoral fellowship through the Yale Cancer Biology Training Program (T32-CA193200) and is supported by an NCI Research Supplement (R01-CA276108-02S2). D.C.M. is supported by the Yale Medical Scientist Training Program (5T32-GM007205). S.S.A. was supported by a postdoctoral fellowship through the Yale Cancer Biology Training Program (T32-CA193200). A.T. was supported by the Yale Science, Technology, and Research Scholars (STARS) II program. R.S.P. acknowledges support from a National Science Foundation Graduate Research Fellowship (NSF-GRFP) and a Ford Foundation Predoctoral Fellowship. A.W. recognizes funding from the Lo Fellowships for Excellence in Stem Cell Research. N.P.M. was supported by the James Hudson Brown-Alexander B. Coxe Fellowship and an American Association for Cancer Research (AACR) Career Development Award. F.S.G. is supported by a Veterans Administration Senior Clinical Scientist Award (BX003250). M.S.R. was supported by R01-DK090489, R01-DK126447, and the Naratil Pioneer Award from the Women’s Health Research at Yale. M.D.M. acknowledges support from an NIH Director’s New Innovator Award (DP2-CA248136), Damon Runyon-Rachleff Innovation Awards (66-21/66S-21) and in part, the Yale Comprehensive Cancer Center Support Grant (P30-CA016359). The content is solely the responsibility of the authors and does not necessarily represent the official views of the National Institutes of Health. This work was supported in large part by a Yale Cancer Center Team Challenge Award (J.W., M.S.R., and M.D.M.), a Lustgarten Foundation Innovation and Collaboration grant (M.D.M.), and NCI R01-CA276108 (M.D.M.).

## AUTHOR CONTRIBUTIONS

C.F.R., J.W., M.S.R., and M.D.M. conceived of and designed the study. C.F.R., X.G., R.M., D.C.M., S.S.A., A.T., M.K., and N.P.M performed tumor studies in mice and cell lines including data acquisition, analysis, and interpretation. J.G., R.S.P., and A.W. developed the diet panel and performed diet studies in wild-type mice, which were supervised by M.S.R. C.F.R. and G.L. performed GC/MS and LC/MS/MS lipidomics analyses on cells and pancreata, which were supervised by D.C. and J.D.H. R.R.V. performed GC/MS lipidomics analyses on diets and plasma, which were supervised by M.C.R. M.E.R. performed pathologic assessment of mouse tumor samples. F.S.G. facilitated acinar cell experiments and interpretation of data. M.D.M. coordinated and supervised the overall study. C.F.R. and M.D.M wrote the manuscript with input from all authors.

## DECLARATION OF INTERESTS

M.D.M. and S.S.A. are inventors on a patent applied for by Yale University that is unrelated to this work. M.D.M. received research funding from a Genentech supported AACR grant and an honorarium from Nested Therapeutics. All other authors declare no competing interests.

## SUPPLEMENTAL INFORMATION

**Table S1. Worldwide food availability and PDAC risk data, Related to Figure 1**. Daily per capita and percent calories macronutrient (fat, protein, carbohydrate) supply for each country worldwide from the Food Agricultural Organization^29^ and age-stratified risk of PDAC (pdac_asr_world) from GLOBOCAN^30^.

**Table S2. Mouse diet panel ingredients, Related to Figure 2**. Fatty acid composition of isocaloric high-fat diets used in the study, categorized by their primary fat source. The table lists the macronutrient composition (protein, carbohydrate, fat) as a percentage of total grams and kilocalories, as well as the specific ingredients and their respective quantities.

**Table S3. Plasma fatty acids in high-fat diet-fed mice, Related to Figure 2**. Average plasma fatty acid concentrations (nmol/mL) measured in non-tumor bearing C57BL/6 mice fed various high-fat diets for 12 weeks. NA indicates values not detected or not measured.

**Table S4. RNA-seq analyses of HFD-fed mice, Related to Figures 3, 5, and 6.** RNA-seq normalized expression counts and differential expression (DEG) analyses of HFD-fed KC (12 weeks and 3 weeks feeding) and KCO mice (8 weeks feeding).

**Table S5. Lipidomics analyses of HFD-fed mice, Related to Figures 3 and 6**. Lipidomics data derived from HFD-fed KC (12 weeks and 3 weeks feeding) and KCO mice (8 weeks feeding). Raw lipid intensity values were first normalized to the amount of inorganic phosphate in nanomoles (normalization_factor) in each sample to account for differences in sample loading (norm_values). The resulting norm_values were then normalized to the response of the internal standard to correct for instrument variability and technical error (normalized_resp).

**Table S6. MRM transitions used for targeted lipidomics analysis, Related to Figures 3 and 6**. Multiple reaction monitoring (MRM) transitions for phosphatidylcholine (PC), lysophosphatidylcholine (LPC), phosphatidylethanolamine (PE), and triglycerides (TG) were measured using an Agilent 6490 QQQ mass spectrometer. The table includes information on the precursor and product ions, dwell time, fragmentor voltage, collision energy, cell accelerator voltage, and polarity for each compound. Internal standards (PC13:0 and PE17:0) were included for normalization and quantification.

## METHODS

### Animal studies

Animal studies were performed at Yale University West Campus and approved under the Yale University Institutional Animal Care and Use Committee (IACUC) protocol #20170. *Kras^LSL-G12D^* (Stock #008179), *Pdx1-Cre* (Stock #014647), *Lep^ob/ob^*(Stock #000632), *MADM11-GT* (Stock #013749), *MADM11-TG* (Stock #013751), *Trp53^KO^* (Stock #002101), and wild-type C57BL/6 (Stock #000664) mice were obtained from the Jackson Laboratory (JAX). KC, KCO, and K-MADM-p53 (*Pdx1-Cre*; *Kras^LSL-G12D^*; *MADM11-GT*/*MADM11-TG*-*Trp53^KO^*) mice were generated by breeding. KC mice were kept on a C57BL/6 background. KCO mice were maintained in a hybrid C57BL/6 and 129/Sv background. K-MADM-p53 mice were of mixed background. PCR genotyping of the mice was performed using GO Taq Green Mastermix (Promega) with template genomic DNA isolated from the tail or ear using Hotshot extraction. Primers and protocols used were previously described^37,70^. For breedings to generate KCO mice, *Lep^ob/ob^* parental mice were administered AAV2/1-CAG-mLeptin via intramuscular injection once, which restores fertility^37^.

HFDs were obtained from Research Diets, Inc., and the complete ingredient list of the isocaloric diets used in the study are detailed in **Table S2**. Given differences in baseline tumor progression kinetics^37^, KC and KCO mice were fed CD or HFDs for 12 or 8 weeks, respectively, starting at 6 weeks of age. For plasma fatty acid analyses or measurements of fasting glucose or insulin, wild-type C57BL/6 male mice were fed safflower or HO safflower diets for 12 weeks. Following an overnight fast, whole blood was collected in the morning via tail nick using heparinized capillary tubes and centrifuged for 8 minutes at 8000 x *g* 4°C. Glucose was measured using a glucometer (Contour Next) and plasma insulin was measured by ELISA (ALPCO) per manufacturer’s instructions. For fatty acid analyses, fasted mice were euthanized by cervical dislocation, and whole blood was collected by cardiac puncture and processed for plasma. K-MADM-p53 mice were fed CD or HFDs for 4 weeks starting at 3 weeks of age. Analyses of fat mass and lean mass were performed by NMR using an Echo MRI whole body composition analyzer (Echo Medical Systems). Mouse weights were monitored weekly using a small animal scale. HFDs were changed once or twice weekly, and average daily food intake was measured for each cage of mice by weighing starting and ending food and dividing by the number of mice in each cage and duration of feeding. Mice of both sexes were used for all experiments as denoted in the figures and legends. Mice were housed in a specific pathogen-free (SPF) facility and kept at room temperature with standard day-night cycles.

### Tissue preparation for histology, immunohistochemistry, and fluorescence imaging

Mice were euthanized by CO2 asphyxiation. Pancreata were isolated, fixed in 10% neutral-buffered formalin (Electronic Microscopy Sciences) overnight, dehydrated in 70% ethanol, and embedded in paraffin by Yale Tissue Pathology Services. Adjacent 5-μm sections were cut and stained with hematoxylin and eosin (H&E), Sirius Red, or subject to immunohistochemistry (IHC) using a ThermoFisher Scientific Autostainer 360. The following primary antibodies were used: mouse anti-Ck19 (DSHB TROMA-III, 1:500), rabbit anti-Ki67 (Biocare Medical CRM325, 1:200), rat anti-p19^ARF^ (Santa Cruz Biotech sc-32748, 1:100), rat anti-Cd45 (Abcam ab25386, 1:100), rabbit anti-F4/80 (CST 70076, 1:400), rabbit anti-SMA (Invitrogen PA1-37024, 1:100), mouse anti-B220 (BD Pharmingen 550286, 1:40), rabbit anti-Cd3 (CST 78588, 1:200), rabbit anti-4-HNE (Abcam, ab46545, 1:200), and rabbit anti-Nqo1 (Sigma-Aldrich HPA007308, 1:100). Mach 2 HRP-labeled micro-polymers (Biocare Medical) were used for primary antibody detection. K-MADM-p53 mice were subject to intracardiac perfusion with PBS followed by 4% paraformaldehyde (PFA; Electronic Microscopy Sciences) in PBS, fixed in 4% PFA overnight at 4°C, cryoprotected in 30% sucrose in PBS at 4°C, embedded in O.C.T. (Tissue-TEK), and stored at -80°C. 10-μm cryosections were obtained using a Leica Cryostat (CM1860). Slides were washed in PBS, stained with DAPI (1:500; ThermoFisher Scientific) in PBS, and imaged with a modified Nikon T2R inverted microscope (MVI), 4x/10x/20x/40x objectives, and a 2.8 MP CoolSNAP Dyno CCD camera (Photometrics).

### Pathologic analysis and quantification

Scanned H&E and IHC sections were digitally analyzed using QuPath v0.5 in a blinded fashion. Disease burden was calculated as the percent area of disease (ADM, PanINs, and PDAC) relative to the total pancreas area using QuPath. Morphologic classification of disease lesions was assessed independently by C.F.R. and an expert gastrointestinal pathologist (M.E.R.) for the KC and KCO models and C.F.R., S.A., and A.T. for the K-MADM-p53 model. For each animal, pancreatic tissue sections were examined by light (KC and KCO) or fluorescence (K-MADM-p53) microscopy in blinded fashion. The percentage of the gland that was occupied by ADM, low-grade PanIN, high-grade PanIN, and PDAC was determined. Pathologic assessment was performed at low and high magnification, using established histologic criteria for mouse pancreatic neoplasia^31,33^. Quantitative assessment of chromogenic IHC staining within (for tumor cell features) and around (for microenvironmental cell features) PanIN lesions was performed using QuPath with automated cell segmentation (nucleus, cytoplasm, cell) and analysis of DAB optical density corrected for background staining to determine positive cells for proteins expressed in the nucleus, cytoplasm, or both.

### RNA extraction, qRT-PCR, and bulk RNA-sequencing

Mouse pancreata were snap frozen in liquid nitrogen and manually pulverized using the BioPulverizer (BioSpec Products 59012MS). RNA was extracted from pulverized mouse tissue (20-40 mg) homogenized via a QIAshredder column (Qiagen) or primary acinar cells using the Qiagen Mini RNeasy kit following the manufacturer’s protocol with DNAse I treatment to remove any genomic DNA contamination. 1 μg of RNA was used to synthesize cDNA using Applied Biosystems™ High-Capacity cDNA Reverse Transcription Kit. cDNA was diluted in nuclease-free water (1:5), and qPCR assays were run on a CFX Opus 384 Real-Time PCR System (Bio-Rad) using SYBR Green (ThermoFisher Scientific) with the following primers: *Scd2*-F 5’- GCATTTGGGAGCCTTGTACG-3’, *Scd2-*R 5’-AGCCGTGCCTTGTATGTTCTG-3’, *Fasn*-F 5’- GGAGGTGGTGATAGCCGGTAT-3’, *Fasn*-R 5’-TGGGTAATCCATAGAGCCCAG-3’, *18S-*F 5’-TAAGTCCCTGCCCTTTGTAACACA-3’, and *18S-*R 5’-GATCCGAGGGCCTCACTAAC- 3’. RNA quality and quantity was assessed using an Agilent Bioanalyzer, and only high-quality RNA samples (RIN>7) were used in RNA-seq analyses. RNA-seq libraries were prepared with polyA selection (Illumina) and sequenced (>25M 100-bp paired-end (2x100) reads on NovaSeq (Illumina)) with the Yale Center for Genome Analysis (YCGA). Low-quality bases and adapter sequences were trimmed using Trim Galore (v0.6.7) and CutAdapt (v3.5). Cleaned reads were aligned to the UCSC mouse genome (mm10) using the STAR aligner (v2.7.9). Differential gene expression analysis was performed using DESeq2 in R, and genes with an adjusted *p*-value < 0.05 were considered differentially expressed. Gene Set Enrichment Analysis (GSEA) and functional overrepresentation analysis were performed using the R package clusterProfiler (v4.12.5). For GSEA, the ranked gene list was analyzed using the GSEA() function, with pathways from Gene Ontology (GO) Biological Process database and a previously published ferroptosis transcriptional signature^52^. Enrichment scores were calculated using 1,000 permutations to estimate significance, and adjusted *p*-values were computed using the Benjamini–Hochberg method. For functional overrepresentation analysis, differentially expressed genes (DEGs) with an adjusted *p*-value < 0.05 and absolute log2 fold-change > 1 were used as input for the enrichGO() function in clusterProfiler. Overrepresented biological processes were identified using the GO Biological Process database.

### Cell culture

KP4 and MiaPaCa-2 PDAC cells were obtained from the Riken Bioresource Research Center Cell Bank and the American Type Culture Collection (ATCC) biorepository, respectively, with authentication by STR profiling by the vendors. KPC (KPC-7307) murine PDAC cells were derived from a C57BL/6 *Pdx1-Cre*; *Kras^LSL-G12D/+^; Trp53^LSL-R172H/+^* mouse, as previously described^37^. Cells tested negative for mycoplasma by PCR performed by the Yale Molecular and Serological Diagnostics Core. All PDAC cells were grown under high glucose DMEM (Corning) supplemented with 10% fetal bovine serum (FBS; ThermoFisher Scientific), penicillin-streptomycin (1%; ThermoFisher Scientific), and L-glutamine (2mM; ThermoFisher Scientific). KP4 knockout clones (KO22 and KO63) were generated by CRISPR/Cas9-mediated knockout, as previously described^60^. MiaPaCa-2 clones resistant to KRAS^G12C^ inhibition (SR5 and SR6) were generated by chronic treatment with 1 µm sotorasib (Selleck Chem) for two months with manual picking of surviving clones. Genomic DNA was extracted using a High-Pure PCR template preparation kit (Roche), and KRAS exons were amplified for amplicon sequencing (MGH CCIB DNA Core) or Sanger sequencing (Yale Keck DNA Sequencing Core) using published primer sets^60^. KPC cells were grown for 72 hours in DMEM supplemented with 10% charcoal-stripped FBS (ThermoFisher Scientific) in the presence of 200 µm OA (Sigma) prior to GC/MS fatty acid analysis. Human PDAC cells were grown in the presence or absence of OA, LA (Sigma), RSL-3 (Selleck Chem), or ML162 (Selleck Chem) for 72 hours. Cell viability was assessed by the CellTiter-Glo (CTG) assay (Promega) with luminescence read using a plater reader (Synergy H1). Primary acinar cells were isolated from 6–8-week-old WT or 9-week-old KC mice (pre-fed with CD or HFDs for 3 weeks), as previously described^81^. Acinar cells were treated with OA, LA, RSL-3, or ML162 for 6 hours prior to RNA isolation (for qRT-PCR analysis) or the CTG assay.

### Western Blot

PDAC whole cell lysates were harvested in RIPA buffer (Pierce), EDTA (1:100) (ThermoFisher Scientific), and Halt protease and phosphatase inhibitor cocktails (1:100) (ThermoFisher Scientific), placed on a rotator at 4°C for 15 min, and centrifuged at maximum speed at 4°C for 20 minutes. Protein concentrations were measured using the BCA Protein Assay Kit (Pierce) following the manufacturer’s protocol. Equal amounts of protein (30 µg) were loaded to each well of a mini-PROTEAN 4-20% TGX stain-free precast gels (Bio-Rad). Protein was transferred to nitrocellulose membranes using the TransBlot Turbo Transfer System (Bio-Rad). Membranes were blocked for one hour using Odyssey Blocking Buffer (Licor) diluted 1:2 in phosphate buffered saline (PBS), then incubated with primary antibodies diluted in Odyssey Blocking Buffer and PBS-0.1% Tween (PBS-T) (1:2) overnight at 4°C. Primary antibodies included mouse anti-KRAS (1:200, clone 3B10-2F2, Sigma-Aldrich) and rabbit anti-HSP90 (1:5000, Cell Signaling Technologies 4877). The specificity of the KRAS antibody was confirmed using RASless mouse embryonic fibroblasts^82^ expressing a single RAS isoform (obtained from the National Cancer Institute (NCI) RAS Initiative). Membranes were washed for 10 minutes with PBS-T three times and then incubated in DyLight secondary antibodies (1:5000, Cell Signaling Technologies) for one hour at room temperature. Membranes were washed with PBS-T for 10 minutes, three times each, prior to imaging using the ChemiDoc Touch Imaging System (Bio-Rad).

### Lipidomics

Fatty acids in diets and mouse plasma from C57BL/6 male mice fed each HFD for 12 weeks were analyzed by gas chromatography mass spectrometry (GC/MS), as previously described^28,83^. Quantitative fatty acid data were normalized to plasma volume or the mass of input diet. Average plasma fatty acid levels are shown in **Table S3**. Pearson correlation analyses of dietary or average plasma fatty acid levels with median disease burden were performed in R using the ggpubr package (v0.6.0). Palmitate levels in KPC cells were measured by GC/MS, as previously described^84^. In brief, cells were harvested in 0.9% NaCl and centrifuged at 10,000 RPM at 4°C. The pellet was resuspended in 2:1 chloroform:methanol. Prior to drying under nitrogen, 50 nmol of heptadecanoic acid was added to all samples as an internal control. Following saponification, metabolites were dried under nitrogen again and methylated with boron trifluoride (Sigma). Mass spectral data were obtained on an Agilent 7890B Gas Chromatograph coupled with an Agilent 5977A MDS. The GC/MS settings were as follows: The injection volume was 1 µL with a split ratio of 2:1. The injector temperature was maintained at 280°C. The column (Agilent 19091S-433UI: HP-5ms Ultra Inert) was operated at a flow rate of 1.2 mL/min with helium as the carrier gas. The oven temperature program was as follows: 50°C for 1 min, ramped to 60°C at 40°C/min and held for 5 min, increased to 270°C at 25°C/min and held for 6 min, increased to 300°C at 10°C/min and held for 6 min, and finally increased to 310°C at 15°C/min and held for 1.4 min. The transfer line temperature was set at 280°C. Electron ionization (EI) was performed at 70 eV, and data were collected in scan mode. Palmitate was monitored at 270-286 m/z.

For liquid chromatography tandem mass spectrometry (LC/MS/MS) analyses, lipids were extracted from pancreatic tissue using a modified Folch method^85,86^. Briefly, frozen pancreatic tissue (∼10-20 mg) was pulverized using a BioPulverizer (BioSpec Products 59012MS) under liquid nitrogen to prevent thawing. Samples were homogenized in a 1:1 mixture of chloroform:methanol (500 µl) containing an internal standard (50 µl, 5000 pmol/mL stock), using ceramic beads (MP #6910050) and an orbital shaker (three 10-second pulses at 5.5 Hz with 5-minute rest intervals). Following centrifugation at 15,000 × g for 5 minutes, the supernatant was transferred to a glass tube and extracted using a modified Folch ratio of 2:2:1.8 (chloroform:methanol:0.9% NaCl). The organic phase was collected, and ∼1/4 of the volume was stored at -80°C for inorganic phosphate analysis, as previously described^87^. The remaining volume was dried under nitrogen at 50°C and reconstituted in 150 µl Zorbax B solution (90% isopropanol, 9% acetonitrile, 1% water) for lipidomic analysis. Pancreatic phospholipids were measured by LC/MS/MS with a slight modification from a previous report^88^. Briefly, phospholipids extracts were spiked with internal standards PC13:0 and PE17:0 (Avanti Polar Lipids). Samples were prepared in silanized 200-µL injection inserts and analyzed on an Agilent 6490 QQQ mass spectrometer with an Agilent 1290 series HPLC system. 3 µL samples were injected into a ZORBAX eclipse plus C18 column (2.1x100mm 1.8mm, Agilent) with the thermostat set at 50°C for separation of molecular species by gradient elution and detected by MS, using scheduled multiple reaction monitoring (MRM) (**Table S6**). In this manner, a peak with unique column elution time and mass-to-fragment profile was measured. Data were analyzed using Agilent MassHunter Quantitative and Quantitative analysis software. The peaks of phospholipid species were integrated and normalized to inorganic phosphate and the internal standards to determine relative abundance.

### Analyses of human data on macronutrient availability, fat consumption, circulating metabolites, and PDAC risk

Macronutrient food availability data for each country were extracted from the Food Agricultural Organization of the World Health Organization (WHO)^29^ in the FAOSTAT database (https://www.fao.org/faostat/en/#home). Age-standardized risk (ASR) of PDAC for each country were derived from the Global Cancer Observatory (GLOBOCAN) 2022 dataset^30^ (https://gco.iarc.fr/today/en). Tabulated data are included in **Table S1**. Spearman correlation analyses were performed in R using the ggpubr package (v0.6.0), excluding countries that did not have both food availability and PDAC risk data. Food availability data for 2014 were used in correlation analyses, given a predicted ∼8-year lag between the initial transformation event and PDAC diagnosis based on genetic evolution studies^89^. Longitudinal data for fatty acid availability and consumption in the United States were extracted from the United States Department of Agriculture (USDA; https://www.ers.usda.gov/data-products/food-availability-per-capita-data-system) and National Health and Nutrition Examination Survey (NHANES; https://data.cdc.gov/NCHS/NHANES-Select-Mean-Dietary-Intake-Estimates/), respectively. Circulating metabolite associations with incident PDAC risk in the UK Biobank cohort were performed using the online research tools for the Nightingale Biomarker-Disease Atlas (https://research.nightingalehealth.com/atlas) to generate forest plots and biomarker-wide association plots. NMR spectroscopy-based biomarker profiling of plasma samples was approved under UK Biobank Project 301418. Methods for NMR spectroscopy-based metabolite measurements in the cohort were previously described^48^. Hazard ratios for biomarkers (corrected for spectrometer) with incident PDAC (based on ICD-10 codes, 1789 cases) were calculated based on Cox regression analysis adjusted for age, sex, and UK biobank assessment center across the full cohort (487,689 individuals) with a significance cut-off of *p* < 0.000005.

### Statistical Analyses

Statistical analyses were performed using R as described in the figure legends. Comparisons between two groups were performed using the Welch test (for normally distributed data) or Wilcoxon rank-sum test (for non-normally distributed data). Multiple group comparisons were performed using one-way ANOVA (for normally distributed data) or Kruskal-Wallis test (for non-normally distributed data) with Tukey’s post-hoc test or Dunn’s post-hoc test to account for multiple comparisons, respectively. Differences in weight gain were analyzed using the repeated measures ANOVA (rmANOVA) followed by post-hoc comparisons using estimated marginal means (EMMeans). Contingency analyses were performed with Fisher’s exact test. Cox regression analysis was used for association of plasma metabolites with PDAC incidence in humans. Pearson (for normally distributed data) and Spearman (for non-normally distributed data) correlation analyses were performed to determine associations between two continuous variables. *p* < 0.05 was used as level of significance for all statistical analyses.

---

**Figure S1.**
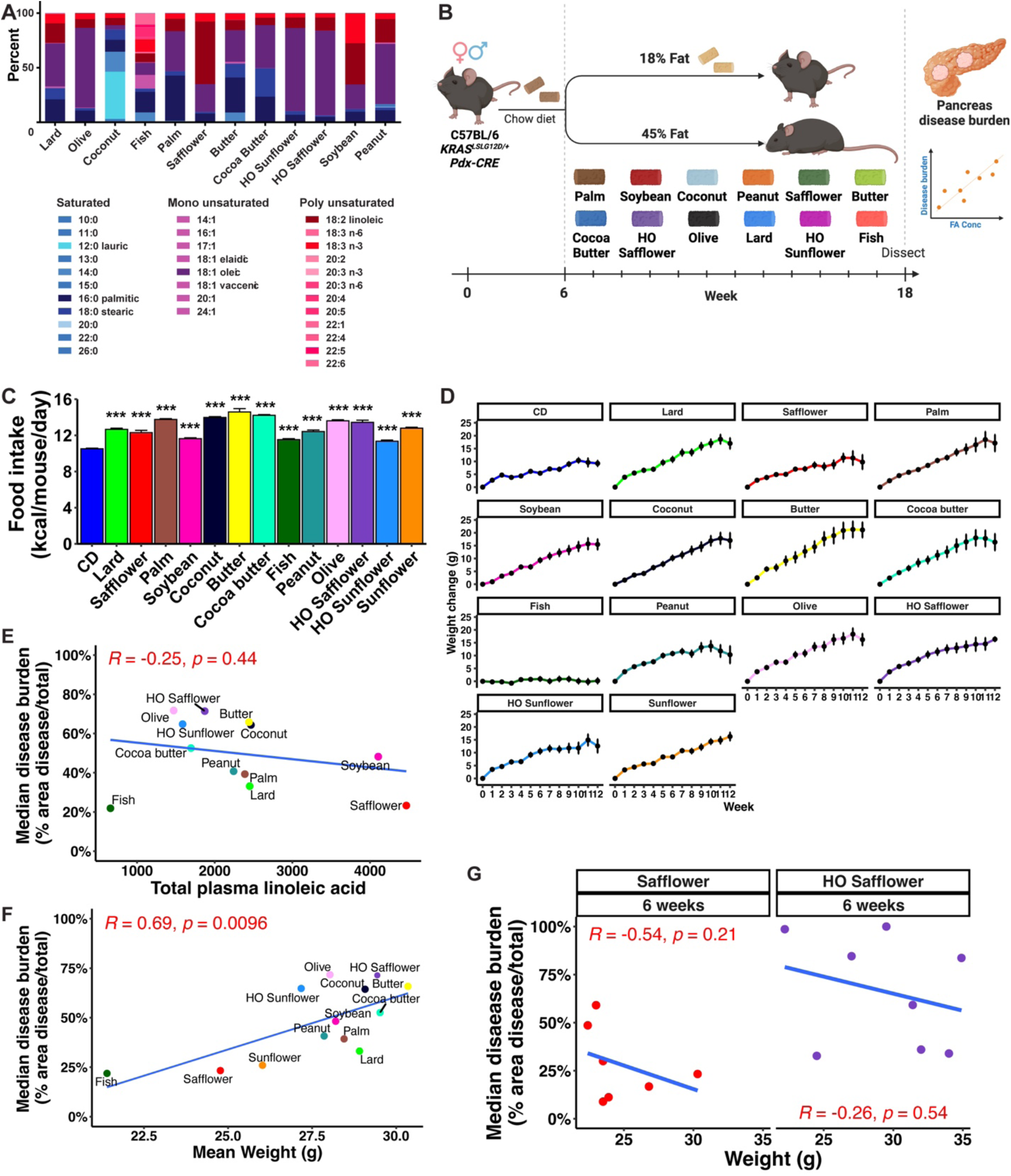
Dietary and physiologic measurements of male mice in HFD screen, Related to Figure 2. (A) Fatty acid composition of isocaloric 45% kcal HFD panel showing the percentage of saturated, monounsaturated, and polyunsaturated fatty acids of each diet by lipidomics analysis. (B) Experimental schema for HFD screen in KC mice randomized into different diet groups. Diets were administered for 12 weeks starting at 6 weeks of age. At the endpoint, the pancreas was harvested for analyses of tumor phenotypes. (C) Food intake (kcal/mouse/day, mean +/- s.e.m., n = 4-13 mice/diet) for male KC mice on each diet in (B). *** p < 0.001, Kruskal-Wallis with Dunn’s post-hoc test. (D) Weight gain (grams (g), mean +/- s.e.m.) for male KC mice in (C). (E) Lack of correlation between median disease burden in male KC mice (n = 4-12 mice/diet) and mean total plasma LA levels (nmol of FA/mL plasma) in non-tumor-bearing C57BL/6 male mice fed each diet for 12 weeks (n = 5 mice/diet). Pearson correlation coefficient (R) and *p*-value are shown. (F) Positive correlation between median disease burden and mean body weight (after 6 weeks HFD feeding, prior to confounding effects of cancer-induced weight loss or malabsorption^90^) of KC mice in (E) across diets. Pearson correlation coefficient (R) and *p*-value are shown. (G) Lack of correlation between disease burden and body weight (after 6 weeks HFD feeding) for mice fed safflower and HO safflower HFDs (n = 7-8 mice/diet). Pearson correlation coefficient (R) and *p*-value are shown.

**Figure S2.**
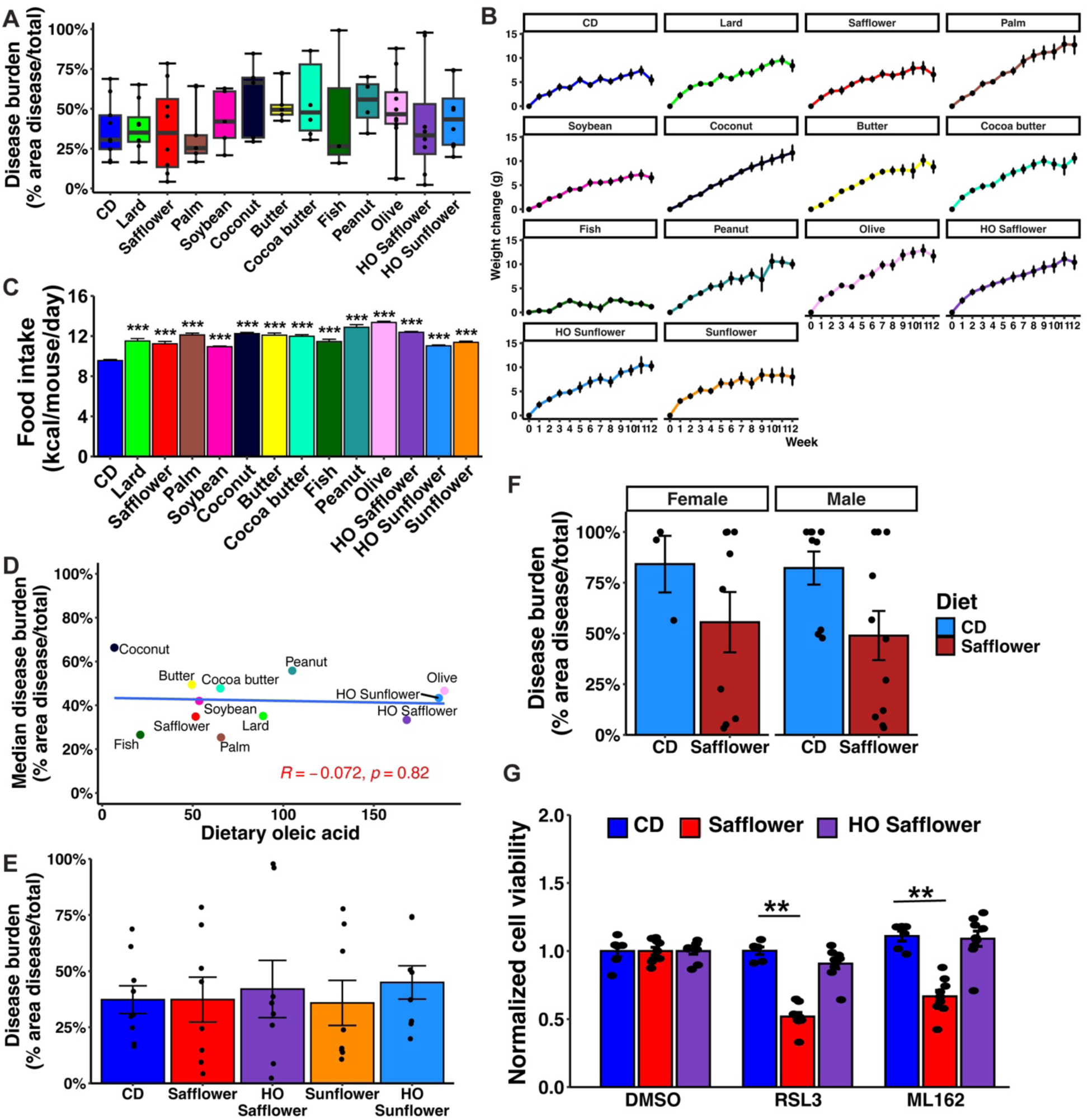
Dietary and physiologic measurements of female mice in HFD screen, Related to Figures 2 and 6. (A) Disease burden (% disease/total pancreas area) for female KC mice fed control diet (CD) or designated HFDs for 12 weeks. Box plots designate 25^th^, 50^th^, and 75^th^ percentiles +/- min/max, n = 3-10 mice/diet. No statistically significant differences across diets were observed, Kruskal-Wallis test with Dunn’s post-hoc test. (B) Weight gain (grams (g), mean +/- s.e.m., n = 5-10 mice per diet) for female KC mice fed each diet for 12 weeks. (C) Food intake (kcal/mouse/day, mean +/- s.e.m.) for female KC mice on each diet in (B). *** p < 0.001, Kruskal-Wallis with Dunn’s post-hoc test. (D) Lack of correlation between median disease burden and dietary oleic acid (µg/mg diet) in female KC mice in (A). Pearson correlation coefficient (R) and *p*-value are shown. (E) Disease burden (mean +/- s.e.m., n = 7-8 mice/diet) for female KC mice fed CD, LODs, or HODs for 12 weeks. No statistically significant differences across diets were observed, Kruskal-Wallis test with Dunn’s post-hoc test. (F) Disease burden (mean +/- s.e.m., n = 3-11 mice/group) for KCO mice on CD or safflower HFD by sex. (G) Cell viability (mean +/- s.d., normalized to vehicle) of primary acinar cells harvested from female KC mice fed CD, safflower HFD, or HO safflower HFD for 3 weeks (n = 2 pooled mice/diet) and treated with GPX4 inhibitors (RSL3 or ML162) vs. vehicle (DMSO) for 6 hours. Dots represent technical replicates (n = 3) from a representative biologic replicate. ** p < 0.01, Kruskal-Wallis test with Dunn’s post-hoc test.

**Figure S3.**
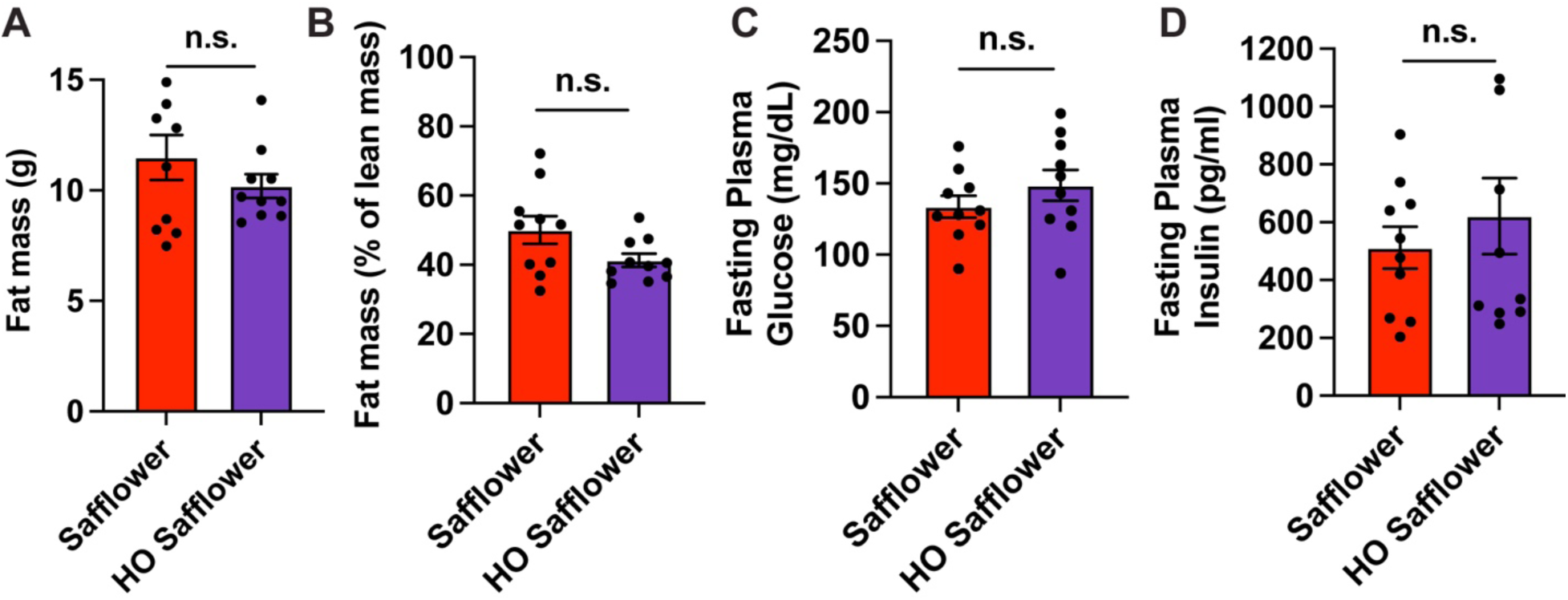
Fat mass and glucose homeostasis measurements in mice fed HFDs differing in oleic acid abundance, Related to Figure 3. (A) Fat mass (grams (g), mean +/- s.e.m., n = 10 mice/diet) in male WT C57BL/6 mice fed safflower (low-oleic) or HO safflower (high-oleic) HFDs for 12 weeks measured by NMR-based Echo MRI. n.s. = non-significant, Welch test. (B) Fat mass (% of lean mass, mean +/- s.e.m.) of mice in (A). (C) Fasting glucose (mean +/- s.e.m.) of mice in (A). (D) Fasting insulin (mean +/- s.e.m.) of mice in (A).

**Figure S4.**
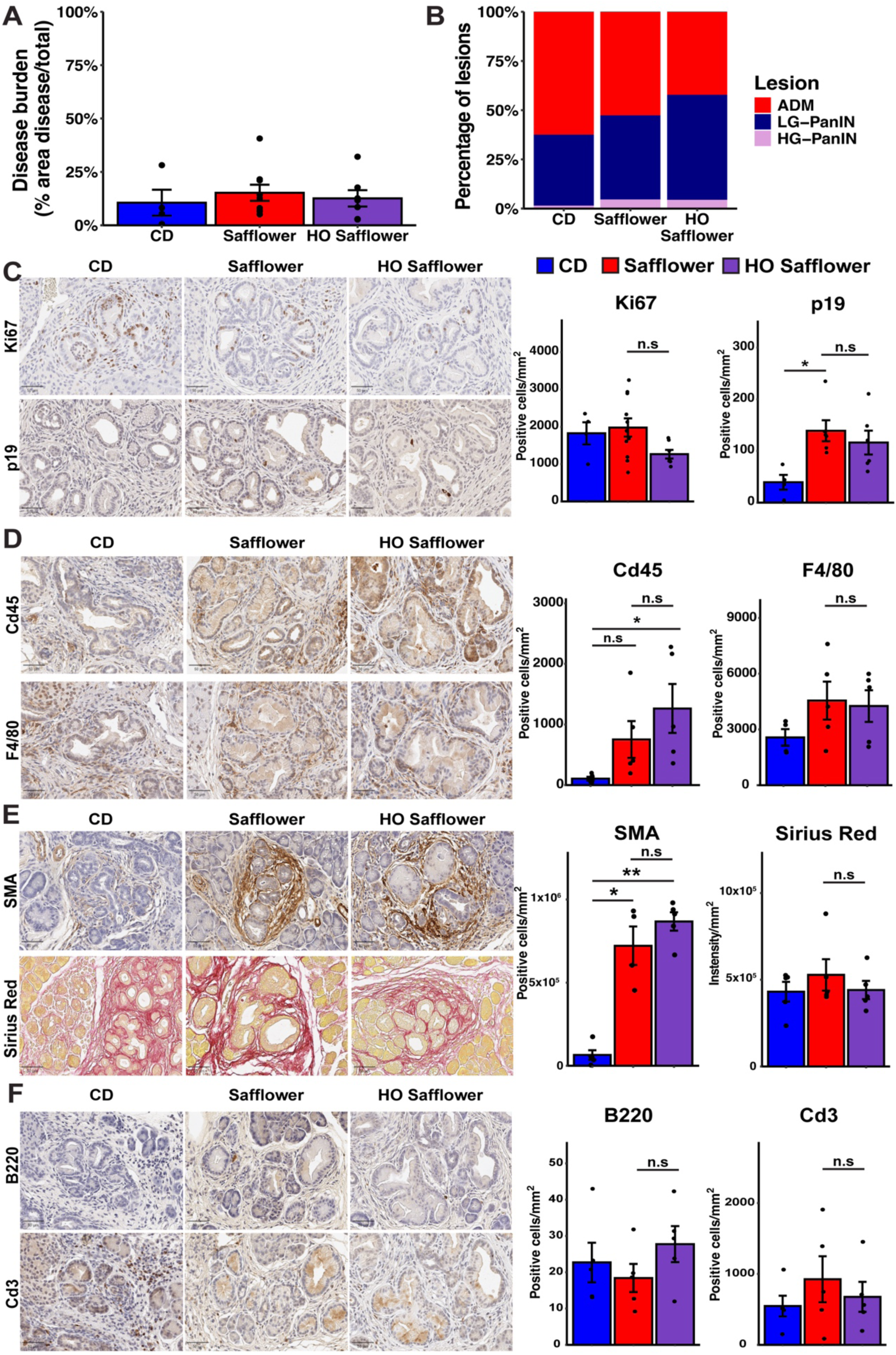
Immunohistochemical analyses of tumor cell and microenvironmental features in HFD-fed mice, Related to Figure 3. (A) Disease burden (mean +/- s.e.m., n = 6-7 mice/diet) for male KC mice fed control diet (CD) or HFDs (safflower, HO safflower) for 3 weeks. No significant differences were observed based on diet, one-way ANOVA. (B) Proportion of pancreatic lesions categorized as acinar-to-ductal metaplasia (ADM), low-grade pancreatic intraepithelial neoplasia (LG-PanIN), and high-grade PanIN (HG-PanIN) in the pancreata of male KC mice in (A). No significant differences were observed between HFDs and CD, Fisher’s exact test. (C) Representative IHC images (left) and quantification (right, mean +/- s.e.m.) of Ki67 and p19ARF immunostaining in PanINs of KC mice fed CD, safflower HFD, and HO safflower HFD for 3 weeks (n = 4-11 mice/diet). * p < 0.05, one-way ANOVA with Tukey’s post-hoc test. n.s. = not significant. Scale bars are 50 µm. (D) Representative IHC images (left) and quantification (right, mean +/- s.e.m.) of pan-leukocyte marker Cd45 and macrophage marker F4/80 immunostaining in PanINs (n = 4-5 mice/diet). * p < 0.05, one-way ANOVA with Tukey’s post-hoc test. n.s. = not significant. Scale bars are 50 µm. (E) Representative IHC images (left) and quantification (right, mean +/- s.e.m.) of myofibroblast marker smooth muscle actin ^28^ immunostaining and collagen staining (Sirius Red) in PanINs (n = 4-5 mice/diet). * p < 0.05, ** p < 0.01, one-way ANOVA with Tukey’s post-hoc test. n.s = not significant. Scale bars are 50 µm. (F) Representative IHC images (left) and quantification (right, mean +/- s.e.m.) of B lymphocyte marker B220 and T lymphocyte marker Cd3 in PanINs (n = 4-5 mice/diet). n.s. = not significant, one-way ANOVA. Scale bars are 50 µm.

**Figure S5.**
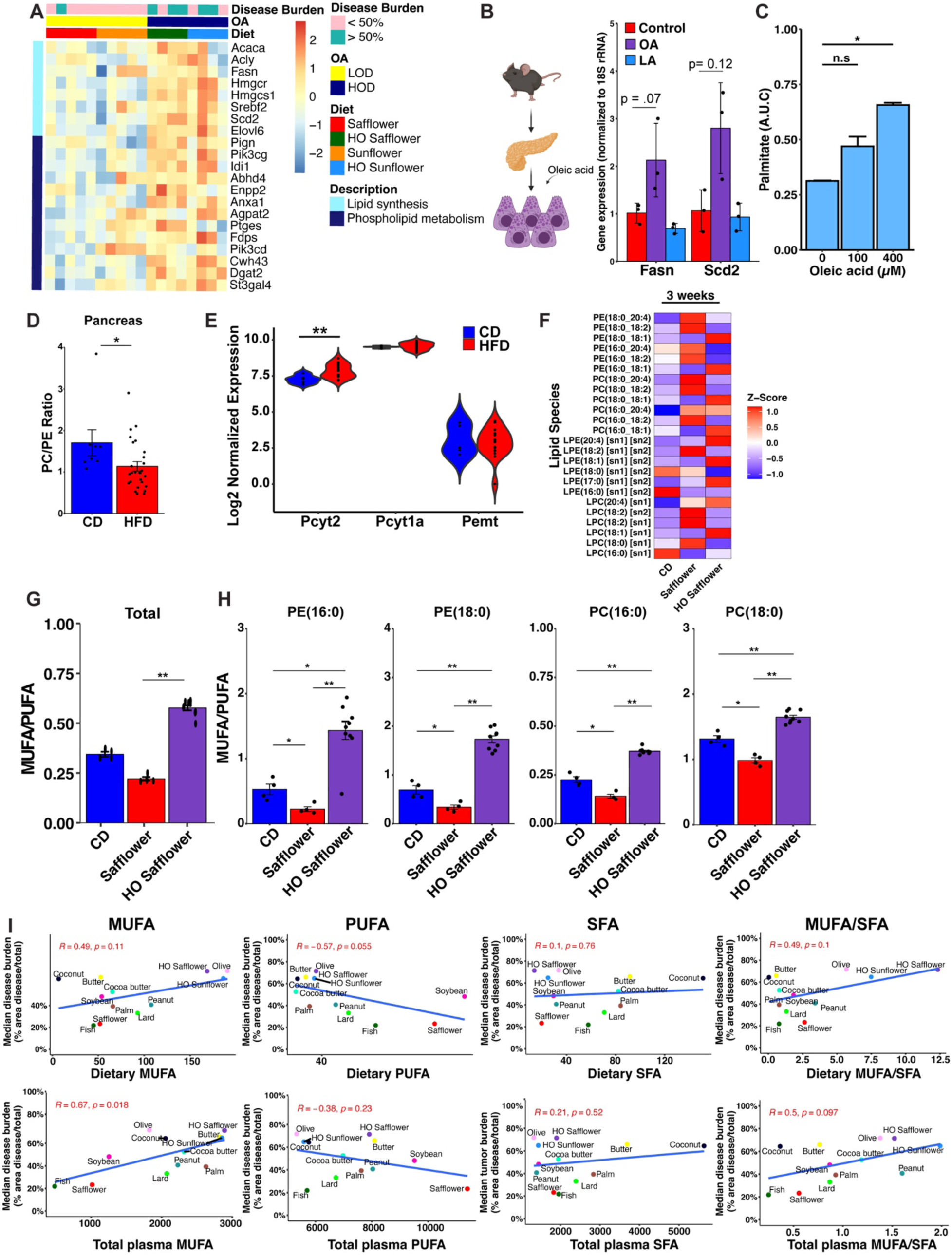
Dietary remodeling of pancreatic lipid metabolism and phospholipid composition, Related to Figure 3. (A) Heatmap of row-normalized expression (RNA-seq) of differentially expressed genes involved in lipid synthesis and phospholipid metabolism (derived from Gene Ontology Biological Process (GO:BP)) in pancreata of LOD- vs. HOD-fed mice (n = 8-10 mice/group). Disease burden is categorized as either <50% or >50% of total pancreas area on histologic sections of adjacent regions. (B) qRT-PCR gene expression (mean +/- s.d., normalized to 18S and to control) of *Fasn* and *Scd2* in primary acinar cells (pooled from n = 2 mice) derived from 6-week-old wild-type C57BL/6 mice and treated with oleic acid (OA, 200 µM) or linoleic acid (LA, 200 µM) for 6 hours *in vitro*. *p*-values were calculated using the Welch test. Dots represent technical replicates (n = 3) from a representative biologic replicate. (C) Relative palmitate abundance (mean +/- s.d.) measured by GC-MS in *KPC* cells (n = 2 biologic replicates) treated with increasing doses of oleic acid (0, 100, 400 µM) over 72 hours. * p < 0.05, Welch test. (D) The ratio (mean +/- s.e.m.) of phosphatidylcholine and phosphatidylethanolamine (PC/PE) in pancreata of male KC mice fed CD (n = 8) or HFDs (safflower, HO safflower, sunflower, or HO sunflower, n = 27) for 12 weeks. * p < 0.05, Welch test. (E) Violin plots of log2 normalized expression counts (denoting 25^th^, 50^th^, and 75^th^ percentiles with min/max) of rate limiting enzymes in PE synthesis (*Pcyt2*), PC synthesis (*Pcyt1a),* and PE to PC conversion (*Pemt*) in RNA-seq analysis of pancreata of KC mice fed CD (n = 5) or HFDs in (**d**) (n = 16) for 12 weeks. ** p < 0.01, Welch test. (F) Heatmap of lipid species detected by targeted LC-MS/MS-based lipidomics in the pancreata of male KC mice fed designated diets for 3 weeks (n = 4-9 mice/diet). Z-scores were generated using relative abundance, and rows are scaled. (G) Total MUFA/PUFA ratio (mean relative abundance +/- s.e.m. of OA(18:1)/LA(18:2) side chains across all PC, PE, LPC, LPE, and TG species measured) of pancreata of male KC mice in (F). ** p < 0.01, Kruskal-Wallis test with Dunn’s post-hoc test. (H) MUFA/PUFA (OA(18:1)/LA(18:2) side chains) ratio (mean relative abundance +/- s.e.m.) in membrane phospholipids (PC and PE) of mice in (F). * p < 0.05, ** p < 0.01, Kruskal-Wallis test with Dunn’s post-hoc test. (I) Correlation of dietary (µg/mg diet) and plasma levels (nmol of FA/mL plasma) for MUFA, PUFA, SFA, and MUFA/SFA (in non-tumor-bearing C57BL/6 male mice fed each diet for 12 weeks, n = 5 mice/diet) with median disease burden in male KC mice (n = 4-12 mice/diet). Pearson correlation coefficient (R) and *p*-value are shown.

**Figure S6.**
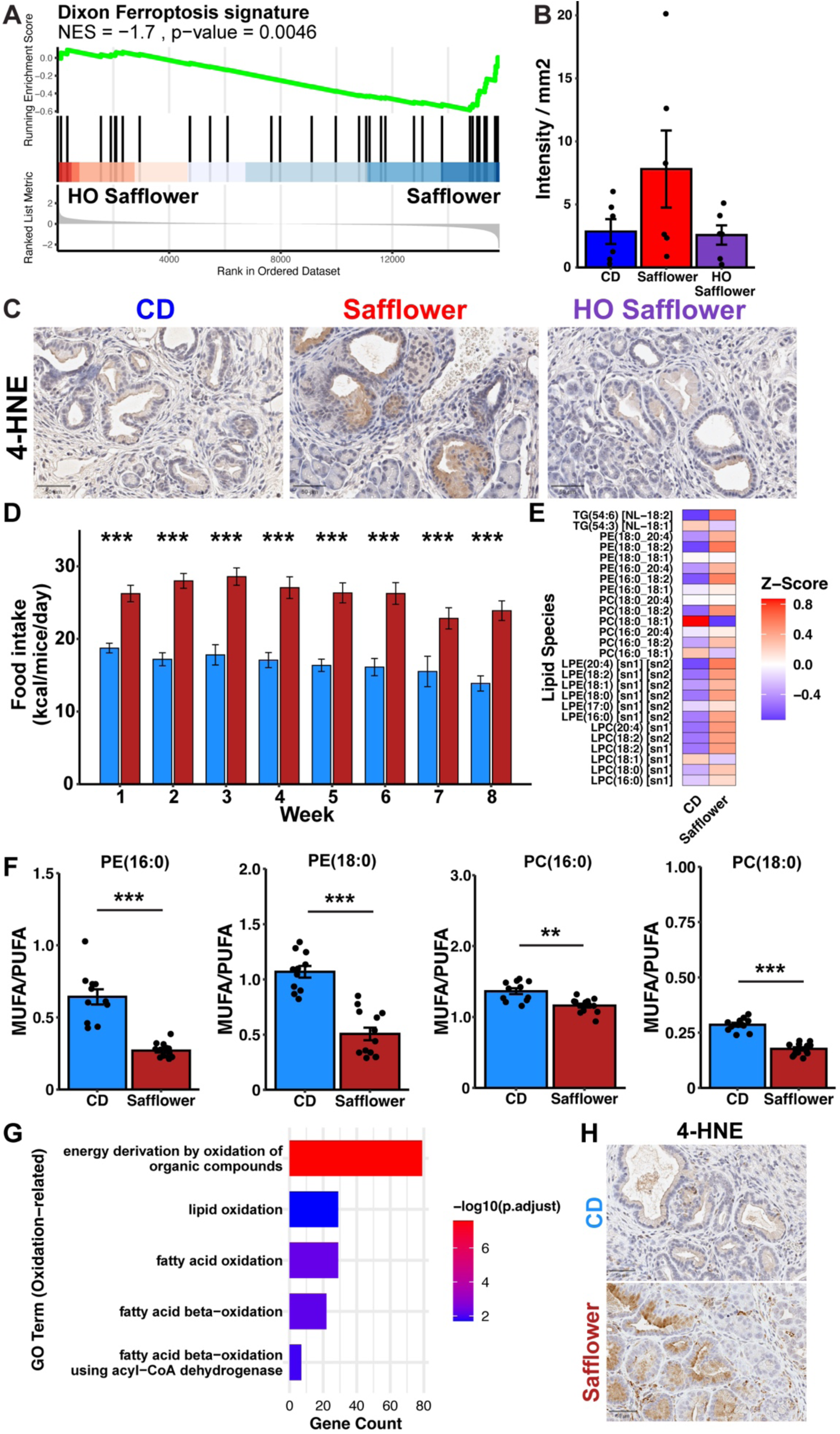
Dietary PUFAs remodel pancreatic phospholipids and induce markers of lipid peroxidation and ferroptosis sensitivity, Related to Figures 5 and 6. (A) GSEA reveals enrichment of the Dixon Ferroptosis Signature (erastin-treated HT-1080 cells^52^) in pancreata of KC mice fed LOD (safflower) vs. HOD (HO safflower) for 3 weeks. Normalized enrichment score (NES) and *p*-value are shown. (B) Quantification (intensity/mm^2^, mean +/- s.e.m., n = 5-6 mice/diet) of 4-hydroxynonenal (4- HNE) immunostaining in PanIN lesions of mice fed the designated diets for 3 weeks. (C) Representative images of 4-HNE immunostaining in (B). Scale bars are 50 µm. (D) Food intake (kcal/day/mouse, mean +/- s.e.m.) over 8 weeks for KCO mice fed control diet (CD) or safflower HFD. * p < 0.05, ** p < 0.01, *** p < 0.001, rmANOVA followed by post-hoc comparisons using EMMeans. (E) Heatmap of lipid species detected by targeted LC-MS/MS-based lipidomics in the pancreata of KCO mice fed designated diets for 8 weeks (n = 11-13 mice/diet). Z-scores were generated using relative abundance, and rows are scaled. (F) MUFA/PUFA (OA(18:1)/LA(18:2) side chains) ratio (mean relative abundance +/- s.e.m.) in membrane phospholipids (PC and PE) of mice in (E). ** p < 0.01, *** p<0.001, Wilcoxon rank-sum test. (G) Functional representation analysis of differentially expressed genes from RNA-seq analyses of pancreata of KCO mice fed safflower HFD for low (<50%) vs. high (>50%) disease burden (n = 3-7 mice/group). Terms related to lipid oxidation (Gene Ontology (GO) Biological Processes) are significantly enriched. (H) Representative IHC image of 4-HNE immunostaining in tumors from KCO mice fed CD or safflower HFD for 8 weeks. Scale bars are 50 µm.

