## Supplementary material for "Diet-induced phospholipid remodeling dictates ferroptosis sensitivity and tumorigenesis in the pancreas": Table S2

### Table S2: Mouse diet panel ingredients

| | D12451 | | D06022403 | | D05122103 | | D03022403 | | D07081501 | | D02062102 | | D06022405 | | D11112703 | | D07062503O | | D05122103 | | D05042003 | | D16010705 | | D21091006 | | D20081706 | |
|---|---|---|---|---|---|---|---|---|---|---|---|---|---|---|---|---|---|---|---|---|---|---|---|---|---|---|---|---|
| **Product #** | **Lard** | | **Olive Oil** | | **Hydro Coconut Oil** | | **Fish (Menhaden) Oil** | | **Palm Oil** | | **Safflower Oil** | | **Butter** | | **Cocoa Butter** | | **Hi Oleic Sunflower** | | **Hi Oleic Safflower** | | **Soybean Oil** | | **Peanut Oil** | | **Sunflower Oil** | | **Control Diet** | |
| % | gm | *kcal* | gm | *kcal* | gm | *kcal* | gm | *kcal* | gm | *kcal* | gm | *kcal* | gm | *kcal* | gm | *kcal* | gm | *kcal* | gm | *kcal* | gm | *kcal* | gm | *kcal* | gm | *kcal* | gm | *kcal* |
| Protein | 23.7 | *20* | 23.7 | *20* | 23.7 | *20* | 23.7 | *20* | 23.7 | *20* | 23.7 | *20* | 23.7 | *20* | 23.7 | *20* | 23.7 | *20* | 23.7 | *20* | 23.7 | *20* | 23.7 | *20* | 23.7 | *20* | 20.1 | *20* |
| Carbohydrate | 41.4 | *35* | 41.4 | *35* | 41.4 | *35* | 41.4 | *35* | 41.4 | *35* | 41.4 | *35* | 41.4 | *35* | 41.4 | *35* | 41.4 | *35* | 41.4 | *35* | 41.4 | *35* | 41.4 | *35* | 41.4 | *35* | 62.1 | *62* |
| Fat | 23.6 | *45* | 23.6 | *45* | 23.6 | *45* | 23.6 | *45* | 23.6 | *45* | 23.6 | *45* | 23.6 | *45* | 23.6 | *45* | 23.6 | *45* | 23.6 | *45* | 23.6 | *45* | 23.6 | *45* | 23.6 | *45* | 8.1 | *18* |
| Total | 88.7 | *100* | 88.7 | *100* | 88.7 | *100* | 88.7 | *100* | 88.7 | *100* | 88.7 | *100* | 88.7 | *100* | 88.7 | *100* | 88.7 | *100* | 88.7 | *100* | 88.7 | *100* | 88.7 | *100* | 88.7 | *100* | 90.4 | *100* |
| kcal/gm | 4.73 | | 4.73 | | 4.73 | | 4.73 | | 4.73 | | 4.73 | | 4.73 | | 4.73 | | 4.73 | | 4.73 | | 4.73 | | 4.73 | | 4.73 | | 4.02 | |
| **Ingredient** | **gm** | ***kcal*** | **gm** | ***kcal*** | **gm** | ***kcal*** | **gm** | ***kcal*** | **gm** | ***kcal*** | **gm** | ***kcal*** | **gm** | ***kcal*** | **gm** | ***kcal*** | **gm** | ***kcal*** | **gm** | ***kcal*** | **gm** | ***kcal*** | **gm** | ***kcal*** | **gm** | ***kcal*** | **gm** | ***kcal*** |
| Casein, 80 Mesh | 200 | *800* | 200 | *800* | 200 | *800* | 200 | *800* | 200 | *800* | 200 | *800* | 200 | *800* | 200 | *800* | 200 | *800* | 200 | *800* | 200 | *800* | 200 | *800* | 200 | *800* | 200 | *800* |
| L-Cystine | 3 | *12* | 3 | *12* | 3 | *12* | 3 | *12* | 3 | *12* | 3 | *12* | 3 | *12* | 3 | *12* | 3 | *12* | 3 | *12* | 3 | *12* | 3 | *12* | 3 | *12* | 3 | *12* |
| Corn Starch | 72.8 | *291* | 72.8 | *291* | 72.8 | *291* | 72.8 | *291* | 72.8 | *291* | 72.8 | *291* | 72.8 | *291* | 72.8 | *291* | 72.8 | *291* | 72.8 | *291* | 72.8 | *291* | 72.8 | *291* | 72.8 | *291* | 369 | *1476* |
| Maltodextrin 10 | 100 | *400* | 100 | *400* | 100 | *400* | 100 | *400* | 100 | *400* | 100 | *400* | 100 | *400* | 100 | *400* | 100 | *400* | 100 | *400* | 100 | *400* | 100 | *400* | 100 | *400* | 75 | *300* |
| Sucrose | 172.8 | *691* | 172.8 | *691* | 172.8 | *691* | 172.8 | *691* | 172.8 | *691* | 172.8 | *691* | 172.8 | *691* | 172.8 | *691* | 172.8 | *691* | 172.8 | *691* | 172.8 | *691* | 172.8 | *691* | 172.8 | *691* | 172.8 | *691* |
| Cellulose, BW200 | 50 | *0* | 50 | *0* | 50 | *0* | 50 | *0* | 50 | *0* | 50 | *0* | 50 | *0* | 50 | *0* | 50 | *0* | 50 | *0* | 50 | *0* | 50 | *0* | 50 | *0* | 50 | *0* |
| Soybean Oil | 25 | *225* | 25 | *225* | 25 | *225* | 25 | *225* | 25 | *225* | 25 | *225* | 25 | *225* | 25 | *225* | 25 | *225* | 25 | *225* | 202.5 | *1823* | 25 | *225* | 25 | *225* | 82 | *738* |
| Lard | 177.5 | *1598* | 0 | *0* | 0 | *0* | 0 | *0* | 0 | *0* | 0 | *0* | 0 | *0* | 0 | *0* | 0 | *0* | 0 | *0* | 0 | *0* | 0 | *0* | 0 | *0* | 0 | *0* |
| Olive Oil | 0 | *0* | 177.5 | *1598* | 0 | *0* | 0 | *0* | 0 | *0* | 0 | *0* | 0 | *0* | 0 | *0* | 0 | *0* | 0 | *0* | 0 | *0* | 0 | *0* | 0 | *0* | 0 | *0* |
| Coconut Oil, Hydrogenated | 0 | *0* | 0 | *0* | 177.5 | *1598* | 0 | *0* | 0 | *0* | 0 | *0* | 0 | *0* | 0 | *0* | 0 | *0* | 0 | *0* | 0 | *0* | 0 | *0* | 0 | *0* | 0 | *0* |
| Menhaden Oil | 0 | *0* | 0 | *0* | 0 | *0* | 177.5 | *1598* | 0 | *0* | 0 | *0* | 0 | *0* | 0 | *0* | 0 | *0* | 0 | *0* | 0 | *0* | 0 | *0* | 0 | *0* | 0 | *0* |
| Palm Oil | 0 | *0* | 0 | *0* | 0 | *0* | 0 | *0* | 177.5 | *1598* | 0 | *0* | 0 | *0* | 0 | *0* | 0 | *0* | 0 | *0* | 0 | *0* | 0 | *0* | 0 | *0* | 0 | *0* |
| Safflower Oil | 0 | *0* | 0 | *0* | 0 | *0* | 0 | *0* | 0 | *0* | 177.5 | *1598* | 0 | *0* | 0 | *0* | 0 | *0* | 0 | *0* | 0 | *0* | 0 | *0* | 0 | *0* | 0 | *0* |
| Butter | 0 | *0* | 0 | *0* | 0 | *0* | 0 | *0* | 0 | *0* | 0 | *0* | 177.5 | *1598* | 0 | *0* | 0 | *0* | 0 | *0* | 0 | *0* | 0 | *0* | 0 | *0* | 0 | *0* |
| Cocoa Butter | 0 | *0* | 0 | *0* | 0 | *0* | 0 | *0* | 0 | *0* | 0 | *0* | 0 | *0* | 177.5 | *1598* | 0 | *0* | 0 | *0* | 0 | *0* | 0 | *0* | 0 | *0* | 0 | *0* |
| High Oleic Sunflower Oil | 0 | *0* | 0 | *0* | 0 | *0* | 0 | *0* | 0 | *0* | 0 | *0* | 0 | *0* | 0 | *0* | 177.5 | *1598* | 0 | *0* | 0 | *0* | 0 | *0* | 0 | *0* | 0 | *0* |
| High Oleic Safflower Oil | 0 | *0* | 0 | *0* | 0 | *0* | 0 | *0* | 0 | *0* | 0 | *0* | 0 | *0* | 0 | *0* | 0 | *0* | 177.5 | *1598* | 0 | *0* | 0 | *0* | 0 | *0* | 0 | *0* |
| Sunflower Oil | 0 | *0* | 0 | *0* | 0 | *0* | 0 | *0* | 0 | *0* | 0 | *0* | 0 | *0* | 0 | *0* | 0 | *0* | 0 | *0* | 0 | *0* | 0 | *0* | 0 | *0* | 0 | *0* |
| High Oleic Sunflower Oil | 0 | *0* | 0 | *0* | 0 | *0* | 0 | *0* | 0 | *0* | 0 | *0* | 0 | *0* | 0 | *0* | 0 | *0* | 0 | *0* | 0 | *0* | 0 | *0* | 0 | *0* | 0 | *0* |
| Peanut Oil | 0 | *0* | 0 | *0* | 0 | *0* | 0 | *0* | 0 | *0* | 0 | *0* | 0 | *0* | 0 | *0* | 0 | *0* | 0 | *0* | 0 | *0* | 177.5 | *1598* | 0 | *0* | 0 | *0* |
| Sunflower Oil | 0 | *0* | 0 | *0* | 0 | *0* | 0 | *0* | 0 | *0* | 0 | *0* | 0 | *0* | 0 | *0* | 0 | *0* | 0 | *0* | 0 | *0* | 0 | *0* | 177.5 | *1598* | 0 | *0* |
| Mineral Mix S10026 | 10 | *0* | 10 | *0* | 10 | *0* | 10 | *0* | 10 | *0* | 10 | *0* | 10 | *0* | 10 | *0* | 10 | *0* | 10 | *0* | 10 | *0* | 10 | *0* | 10 | *0* | 10 | *0* |
| DiCalcium Phosphate | 13 | *0* | 13 | *0* | 13 | *0* | 13 | *0* | 13 | *0* | 13 | *0* | 13 | *0* | 13 | *0* | 13 | *0* | 13 | *0* | 13 | *0* | 13 | *0* | 13 | *0* | 13 | *0* |
| Calcium Carbonate | 5.5 | *0* | 5.5 | *0* | 5.5 | *0* | 5.5 | *0* | 5.5 | *0* | 5.5 | *0* | 5.5 | *0* | 5.5 | *0* | 5.5 | *0* | 5.5 | *0* | 5.5 | *0* | 5.5 | *0* | 5.5 | *0* | 5.5 | *0* |
| Potassium Citrate, 1 H2O | 16.5 | *0* | 16.5 | *0* | 16.5 | *0* | 16.5 | *0* | 16.5 | *0* | 16.5 | *0* | 16.5 | *0* | 16.5 | *0* | 16.5 | *0* | 16.5 | *0* | 16.5 | *0* | 16.5 | *0* | 16.5 | *0* | 16.5 | *0* |
| Vitamin Mix V10001 | 10 | *40* | 10 | *40* | 10 | *40* | 10 | *40* | 10 | *40* | 10 | *40* | 10 | *40* | 10 | *40* | 10 | *40* | 10 | *40* | 10 | *40* | 10 | *40* | 10 | *40* | 10 | *40* |
| Choline Bitartrate | 2 | *0* | 2 | *0* | 2 | *0* | 2 | *0* | 2 | *0* | 2 | *0* | 2 | *0* | 2 | *0* | 2 | *0* | 2 | *0* | 2 | *0* | 2 | *0* | 2 | *0* | 2 | *0* |
| FD&C Yellow Dye #5 | 0 | *0* | 0 | *0* | 0.025 | *0* | 0.025 | *0* | 0.025 | *0* | 0.025 | *0* | 0.025 | *0* | 0.025 | *0* | 0.025 | *0* | 0 | *0* | 0.025 | *0* | 0 | *0* | 0.05 | *0* | 0 | *0* |
| FD&C Red Dye #40 | 0.05 | *0* | 0 | *0* | 0.025 | *0* | 0 | *0* | 0 | *0* | 0 | *0* | 0 | *0* | 0 | *0* | 0.025 | *0* | 0.025 | *0* | 0.025 | *0* | 0.04 | *0* | 0 | *0* | 0 | *0* |
| FD&C Blue Dye #1 | 0 | *0* | 0.05 | *0* | 0 | *0* | 0.025 | *0* | 0.025 | *0* | 0.025 | *0* | 0.025 | *0* | 0.025 | *0* | 0 | *0* | 0.025 | *0* | 0 | *0* | 0.01 | *0* | 0 | *0* | 0 | *0* |
| **Total** | **858.15** | ***4057*** | **858.15** | ***4057*** | **858.15** | ***4057*** | **858.15** | ***4057*** | **858.15** | ***4057*** | **858.15** | ***4057*** | **858.15** | ***4057*** | **858.15** | ***4057*** | **858.15** | ***4057*** | **858.15** | ***4057*** | **858.15** | ***4057*** | **858.15** | ***4057*** | **858.15** | ***4057*** | **1008.8** | ***4057*** |
