## Supplementary material for "Diet-induced phospholipid remodeling dictates ferroptosis sensitivity and tumorigenesis in the pancreas": Table S3

**Table S3. Plasma fatty acids in high-fat diet-fed mice.**

| Fatty_acid | Lard | Olive | Coconut | Fish | Palm | Safflower | Butter | Cocoa butter | HO Sunflower | HO Safflower | Soybean | Peanut |
|---|---|---|---|---|---|---|---|---|---|---|---|---|
| **10_0** | 184.31 | 211.68 | 292.66 | 158.00 | 218.83 | 109.72 | 264.45 | 145.07 | 115.91 | 126.12 | 9.31 | 10.71 |
| **12_0** | 191.29 | 188.26 | 1734.34 | 90.70 | 102.41 | 38.74 | 229.36 | 71.31 | 115.88 | 140.96 | NA | 3.52 |
| **14_0** | 88.85 | 38.08 | 1029.95 | 326.67 | 92.11 | 45.56 | 467.52 | 58.65 | 47.59 | 62.88 | 6.46 | 2.83 |
| **14_1** | 4.05 | 2.26 | 33.20 | 15.55 | 5.68 | 3.42 | 32.83 | 4.87 | 3.07 | 2.66 | 1.09 | 1.41 |
| **16_0** | 1454.61 | 737.58 | 1914.05 | 990.34 | 2005.96 | 1086.03 | 2046.01 | 1133.07 | 853.01 | 1085.29 | 1049.64 | 1024.76 |
| **16_1_n_7** | 254.97 | 192.14 | 716.59 | 422.00 | 354.94 | 113.04 | 674.75 | 287.23 | 157.22 | 178.37 | 129.00 | 163.34 |
| **16_2** | 4.54 | 3.19 | 6.62 | 25.76 | 4.38 | 5.74 | 8.86 | 3.45 | 3.11 | 3.94 | 5.42 | 3.41 |
| **18_0** | 470.86 | 148.69 | 649.95 | 337.35 | 492.28 | 534.04 | 645.41 | 594.71 | 332.35 | 538.09 | 389.31 | 359.98 |
| **18_1_n_9** | 2021.63 | 1800.38 | 1936.83 | 418.80 | 2520.91 | 964.20 | 2680.06 | 2243.53 | 2770.94 | 2842.80 | 1241.03 | 2172.02 |
| **18_2_n_6** | 2450.60 | 1471.20 | 2466.80 | 657.17 | 2388.04 | 4467.53 | 2441.33 | 1694.80 | 1586.45 | 1871.93 | 4105.87 | 2243.19 |
| **18_3_n_3** | 317.28 | 162.64 | 368.33 | 358.80 | 259.52 | 218.62 | 335.61 | 216.69 | 163.40 | 162.25 | 818.20 | 181.06 |
| **18_3_n_6** | 154.03 | 103.44 | 160.30 | 80.25 | 213.02 | 209.75 | 176.77 | 126.05 | 118.57 | 128.64 | 158.29 | 151.94 |
| **20_0** | NA | NA | 3.82 | 62.80 | 64.37 | 15.29 | 53.60 | 15.86 | NA | NA | NA | NA |
| **20_1_n_9** | 46.82 | 22.57 | 83.76 | 48.53 | 93.07 | 46.60 | 97.11 | 56.52 | 61.42 | 46.69 | 12.86 | 55.61 |
| **20_2_n_6** | 24.40 | 4.45 | 11.08 | 4.85 | 18.53 | 23.50 | 17.92 | 14.53 | 11.65 | 6.89 | 10.88 | 9.44 |
| **20_3_n_6** | 126.60 | 132.46 | 158.20 | 36.99 | 178.11 | 169.04 | 248.82 | 171.08 | 121.97 | 148.82 | 131.59 | 123.92 |
| **20_3_n_3** | 3.74 | 2.73 | 4.38 | 7.65 | 113.11 | 4.42 | 86.75 | 122.73 | 102.38 | 4.65 | 4.23 | 2.81 |
| **20_4_n_6** | 3271.13 | 3062.45 | 2121.14 | 1443.13 | 3875.65 | 5740.71 | 3964.54 | 4031.76 | 3135.04 | 4995.05 | 3554.44 | 4838.81 |
| **20_5_n_3** | 76.37 | 76.41 | 128.57 | 1948.03 | 95.78 | 64.21 | 190.25 | 101.30 | 52.53 | 124.46 | 155.75 | 69.64 |
| **22_0** | NA | NA | 61.43 | 5.66 | 9.51 | NA | 49.26 | NA | NA | NA | NA | NA |
| **22_1** | NA | NA | 1.93 | NA | 17.76 | NA | 22.60 | 24.89 | 22.65 | NA | NA | NA |
| **22_2_n_6** | 1.07 | NA | 0.23 | 3.05 | 1.93 | 4.25 | 1.57 | 0.95 | 0.75 | 0.76 | 0.85 | 1.37 |
| **22_3** | 0.86 | 0.58 | 1.46 | 0.19 | 3.75 | 3.09 | 2.56 | 3.49 | 3.33 | 4.09 | 1.38 | 2.49 |
| **22_4_n_6** | 14.90 | 3.83 | 11.48 | 9.97 | 11.25 | 43.52 | 12.86 | 8.82 | 6.58 | 15.12 | 11.22 | 20.99 |
| **22_4_n_3** | 1.52 | NA | NA | 1.70 | 11.16 | NA | 9.17 | 5.63 | 9.32 | NA | 1.70 | 1.12 |
| **22_5_n_6** | 28.76 | 23.54 | 35.97 | 45.54 | 43.17 | 97.49 | 42.14 | 41.52 | 21.78 | 66.76 | 23.70 | 64.90 |
| **22_5_n_3** | 15.57 | 10.31 | 27.31 | 103.65 | 34.19 | 16.37 | 34.26 | 23.37 | 19.17 | 13.20 | 30.53 | 14.94 |
| **22_6_n_3** | 237.27 | 276.67 | 262.24 | 975.43 | 378.44 | 325.66 | 529.77 | 425.41 | 257.79 | 364.89 | 501.86 | 303.13 |
